# A tachykinin precursor 1 medullary circuit promoting rhythmic breathing

**DOI:** 10.1101/2023.01.13.523897

**Authors:** Jean-Philippe Rousseau, Andreea Furdui, Carolina da Silveira Scarpellini, Richard L. Horner, Gaspard Montandon

## Abstract

Rhythmic breathing is generated by neural circuits located in the brainstem. At its core is the preBötzinger Complex (preBötC), a region of the medulla, necessary for the generation of rhythmic breathing in mammals. The preBötC is comprised of various neuronal populations expressing neurokinin-1 receptors, the cognate G-protein-coupled receptor of the neuropeptide substance P (encoded by the tachykinin precursor 1 or *Tac1*). Neurokinin-1 receptors are highly expressed in the preBötC and destruction or deletion of neurokinin-1 receptor-expressing preBötC neurons severely impairs rhythmic breathing. Application of substance P to the preBötC stimulates breathing in rodents, however substance P is often associated with nociception and locomotion in various brain regions, suggesting that *Tac1* neurons found in the preBötC may have diverse functional roles. Here, we aim to characterize the role of *Tac1*-expressing preBötC neurons in the generation of rhythmic breathing *in vivo*, as well as motor behaviors. Using a cre-lox recombination approach, we injected adeno-associated virus containing the excitatory channelrhodopsin-2 ChETA in the preBötC region of *Tac1*-cre mice. Using a combination of histological, optogenetics, respiratory, and behavioral assays, we defined the identity and the role of *Tac1* preBötC neurons. These neurons are glutamatergic and their stimulation promotes rhythmic breathing in both anesthetized and freely moving/awake animals, but also triggers locomotion and overcomes respiratory depression by opioid drugs. Overall, our study identifies a new population of excitatory preBötC with major role in rhythmic breathing and behaviors.

## Introduction

Motor rhythms are fundamental for many biological functions including locomotion and breathing. Breathing relies on motor circuits to promote gas exchange and maintain life. Although these circuits are critical for life, they can be interrupted by behaviors such as pain response and vocalization. At the core of the respiratory network is the preBötzinger Complex (preBötC), a collection of neurons essential to produce and sustain breathing (Smith *et al*., 1991). The preBötC region is comprised of diverse populations of neurons encompassing inhibitory and excitatory neurons. Glutamatergic excitatory neurons form roughly half of preBötC neurons with critical roles in the generation of inspiration. A subpopulation of glutamatergic neurons expresses the neuropeptide substance P (encoded by the tachykinin precursor 1 or *Tac1* gene). Substance P in the preBötC stimulates breathing through activation of its cognate neurokinin-1 receptors (NK-1R) (Montandon *et al*., 2016a), and destruction of NK-1R-expressing neurons abolishes breathing (Gray *et al*., 2001). The biological function of preBötC *Tac1* (substance P)-expressing neurons in the regulation of breathing is not known *in vivo*.

Substance P plays a pivotal role in rhythmic breathing, but it is also a key-molecule involved in nociception (Mantyh, 2002), locomotion (Farrell *et al*., 2021), and arousal (Reitz *et al*., 2021). Substance P is released by nociceptive stimuli (Mantyh, 2002), is expressed in circuits regulating nociception such as the dorsal horn of the spinal cord (Chang *et al*., 2019) and the rostral ventromedial medulla, and modulates descending pain circuits (Khasabov *et al*., 2017). Descending *Tac1* circuits in the brainstem mediate behavioral responses, such as the fight-or-flight response, associated with brisk locomotor activity (Barik *et al*., 2018; Kuwaki, 2021). To anticipate the body’s metabolic demand in the event of locomotor nocifensive response, nociceptive stimuli elicit cardio-respiratory responses, such as increased heart rate and augmented breathing (Jafari *et al*., 2017). *Tac1*-expressing medullary circuits involved in nociception or breathing share similar properties: they are sensitive to opioid drugs and expressed neurokinin-1 receptors. Here, we aim to identify the role of *Tac1*-expressing preBötC cells in regulating breathing and motor behaviors, which constitute different components of nocifensive behaviors.

By combining optogenetics, respiratory, and locomotion assays, we determined the role of *Tac1* preBötC cells in regulating rhythmic breathing and locomotion. Using photostimulation, we first showed that *Tac1* preBötC neurons, a subpopulation of glutamatergic neurons, increased respiratory rhythm by promoting inspiration or reducing expiration in both anesthetized and freely behaving mice. In freely moving mice, photostimulation was mostly effective when mice were in calm, but not active, state. Interestingly, stimulation of *Tac1* preBötC cells directly elicited a strong locomotor response suggesting that *Tac1* preBötC cells play a dual role in promoting breathing and locomotion. Rhythmic breathing is dampened by opioid drugs and preBötC cells mediate a major component of respiratory depression by opioid drugs. Because most *Tac1* preBötC cells co-expressed µ-opioid receptors (encoded by the gene *Oprm1*) in the preBötC, their stimulation reversed the effects of opioid drugs on breathing, suggesting that *Tac1* preBötC neurons constitute a robust excitatory neural circuit involved in breathing that promotes breathing during calm states and that can overcome respiratory depression with narcotics.

## Methods

### Animal care, drug and virus acquisition

All procedures were carried out in accordance with the recommendations of the Canadian Council on Animal Care and were approved by St. Michael’s Hospital animal care committee. Experiments were performed on 63 adult mice of either sex, aged between 3-4 months old and weighing between 20g and 40g. Animals were kept on a 12 h light–dark cycle with unrestricted food and water and all experiments were performed during the day. Vglut2-ires-cre (STOCK Slc17a6^tm2(cre)Lowl^/J; Strain # 016963) and *Tac1*-IRES2-Cre-D (B6;129S-*Tac1*^tm1.1(cre)Hze^/J; Strain # 021877) breeders were obtained from The Jackson Laboratory (600 Main Street, Bar Harbor, ME USA 04609). Litters generated were separate and used randomly by the investigators. An adeno-associated virus (AAV_5_-EF1a-DIO-ChETA-eYFP) that expresses ChR2-E123T Accelerated (ChETA) and the fluorescent protein eYFP was purchased from University of North Carolina (UNC vector core, Chapel Hill, NC, USA). The same serotype virus that expresses eYFP but lacks expression of the light-sensitive protein (AAV-EF1a-DIO-eYFP-WPRE-pA; UNC vector core, Chapel Hill, NC, USA) was also used at the same concentration for control experiments. Fentanyl citrate (50 μg/mL, Sandoz) was used with Health Canada exemption and obtained from the St. Michael’s Hospital In-Patient Pharmacy.

### Surgical procedures

All surgeries were performed using standard aseptic techniques. Before surgery, mice were given pain medication: Anafen (5mg/Kg), Dexamethazone (5mg/Kg) and saline for a total volume of 1 ml subcutaneously. Mice were anaesthetized with isoflurane (2-3% in 100% oxygen) and placed in the prone position in a stereotaxic apparatus (model 940, KOPF instruments, Tujunga, CA, USA) with blunt ear bars where anaesthesia was maintained through a nose cone. Adequate depth of anesthesia was determined via beathing and reflex responses to toe pinch and adjusted if necessary. Animal temperature was monitored through a rectal probe and was maintained at 36.5 °C via a heating pad (Kent Scientific corporation, Torrington, CT, USA). An incision was performed in the skin to expose the dorsal skull, which was then levelled horizontally between bregma and lambda. A small craniotomy was made to access the preBötzinger Complex. A needle containing the adeno-associated virus (AAV5-EF1a-DIO-ChETA-eYFP) was slowly lowered in the preBötC 6.8 mm posterior, 1.2 mm lateral and 6.4 mm ventral to bregma. Coordinates were chosen based on the Mouse Brain in Stereotaxic Coordinates (3^rd^ Edition, Paxinos and Franklin) combined with preliminary mapping of viral injections from our team. Needle was lowered with an infusion rate of 80 nl/min provided by a programmable syringe pump (Harvard Apparatus, Holliston, MA, USA) to avoid any clogging. Once in the targeted region, virus was infused at 50 nl/min reaching a total volume of 300 nl. Control animals were injected with a sham virus that lacks expression of the light-sensitive protein (AAV-EF1a-DIO-eYFP-WPRE-pA). Following infusion, needle was maintained in position for 10 minutes before it was slowly removed and skin was sutured over the skull. Position of viral injection was also confirmed through *in situ* hybridization for expression of *Nk1r* (gene coding for NK-1R) and *Eyfp* (gene for eYFP).

For unanesthetized / freely behaving experiments, an optic fiber was fixed over the preBötC following virus injection. Before the skin was sutured, two sterile stainless-steel screws (P1 Technologies, Roanoke, VA, USA) were implanted in the bone, one near the bregma and one near the preBötC craniotomy. A custom-made sterile cannula containing the optical fiber (200μm, 0.5 NA; FP200ERT, Thor Labs, New Jersey, USA) was then positioned on top of the preBötC 6.8 mm posterior, 1.2 mm lateral and 5.5 mm ventral to bregma. The cannula was then fixed in position resting on top of the skull using dental cement (Co-oral-ite dental MFG. CO., Diamond Spring, CA, USA). Cement was spread on both stainless-steel screws to firmly hold the cannula and skin was later sutured over it. In the 3 days post-surgery, mice were given pain medication: Anafen (5mg/Kg), Dexamethazone (5mg/Kg) and saline for a total volume of 1 ml subcutaneously. Animals were maintained 2-4 weeks before functional experiments were performed so virus could be expressed in targeted region.

### Optogenetic stimulation in anesthetized mice

Optogenetic stimulation was used to selectively activate tachykinin precursor 1 (*Tac1*) and vesicular-glutamate transporter 2 (*Vglut2*) expressing cells in the preBötC. Three *Tac1*-IRES2-Cre-D and two *Vglut2*-ires-cre (3 males and 2 females) received the sham virus and served as controls while twenty *Tac1*-IRES2-Cre-D (14 males and 6 females) and five Vglut2-ires-cre (3 males and 2 females) received the AAV5-EF1a-DIO-ChETA-eYFP virus. Two or 4 weeks after surgery, mice were anesthetized with 2-2.5% isoflurane and were spontaneously breathing (50% oxygen gas mixture, balance nitrogen). Adequate depth of anesthesia was determined via beathing and reflex responses to toe pinch and adjusted if necessary. Animal temperature was monitored through a rectal probe and was maintained at 36.5 °C via a heating pad (Kent Scientific corporation, Torrington, CT, USA). Diaphragm muscle activity was recorded using bipolar electrodes sutured to the upper right abdominal wall adjacent to the diaphragm. Electromyography signals were amplified (AM Systems Model 1700, Sequim, Washington, USA), band-pass filtered (300-5000 Hz), integrated and digitized at a sampling rate of 1,000 Hz using PowerLab 4/26 acquisition system and LabChart Version 8 (ADInstruments, Colorado Spring, CO, USA).

Mice were placed in the prone position in a stereotaxic frame (model 940, KOPF instruments, Tujunga, CA, USA) with blunt ear bars where anaesthesia was maintained through a nose cone. As this group did not have a fixed optical fiber over the preBötC, incision was again performed in the skin to expose the bregma and lambda. The dorsal skull was levelled horizontally between bregma and lambda and a small craniotomy was made to access the site of virus injection in the preBötC. An optical fibre (200 μm, 0.5 NA; CFMC52L10, Thor Labs, New Jersey, USA), connected to a laser source (Laserglow technologies, North York, ON, Canada), was then lowered on top of the preBötC 6.8 mm posterior, 1.2 mm lateral and 5.5 mm ventral to bregma. Position was confirmed through continuous high frequency laser stimulation at 10 mW (50 pulses of 20 ms pulses repeated at 20 Hz). Once breathing was stable (~10 min; average breathing rhythm = 20-30 breaths/min), laser stimulation (blue, wavelength = 473 nm) was performed using various settings under control conditions. Settings included 20 ms pulses repeated 10 times at frequencies of 5, 10, 20, 30 and 40 Hz and power of 1, 3, 5 and 10 mW. For all stimulations, each train of 10 pulses were repeated 5 times with one second pause between them.

Following stimulation phases under control conditions, diaphragm activity was recorded for 10 minutes as a baseline sequence before mice were getting an intramuscular injection of the μ-opioid receptor agonist fentanyl (5μg/Kg). 5 minutes were then given for fentanyl to elicit a respiratory rate depression and laser stimulation was once again performed using various settings. Settings included 20 ms pulses repeated 10 times with a frequency of 20, 30 and 40 Hz and power of 10 mW. For all stimulations, each train of 10 pulses had one second pause between them and were repeated over 1 minute. Mice were then euthanized with intracardiac injection of T-61 and brain was harvested for post-mortem histology.

Data was extracted from LabChart Version 8 (ADInstruments, Colorado Spring, CO, USA) and exported to Microsoft Excel and GraphPad Prism 9 (Version 9.3.1; Graphpad Software) for analysis. For each laser stimulation, respiratory rate (breaths/min) and amplitude (volts) as well as inspiratory and expiratory durations were obtained from rectified diaphragm EMG recordings, averaged over the whole stimulation period, and normalized according to the preceding baseline period (rate and amplitude; average of 1 min) or the average of the 5 preceding breathing cycles (inspiratory and expiratory durations). Since respiratory rate, amplitude and inspiratory / expiratory durations varied from one animal to another, these parameters were expressed as percentage change of the values of the preceding baseline period. Figures with absolute values and means are provided in Supplementary Materials. Timing of the laser stimulation inside the respiratory cycle was also determined to measure the induced period. To do so, we calculated the time between the initiation of inspiration and the start of laser stimulation and normalized it according to the preceding respiratory cycle.

### Optogenetic stimulation in freely behaving mice

Respiratory variables in freely behaving mice were recorded using whole body, flow-through plethysmography (Buxco Electronics, DSI, New Brighton, Minnesota, USA). Two *Tac1*-IRES2-Cre-D and Two Vglut2-ires-cre (3 males and 1 females) received the sham virus and served as controls while 10 *Tac1*-IRES2-Cre-D (7 males and 3 females) received the AAV5-EF1a-DIO-ChETA-eYFP virus. Plethysmography chambers measured 21.5cm in diameter and allowed enough room for mice to move freely during physiological recordings. Chambers were continuously ventilated using room air at a rate of 0.9 L/min (normoxia; F_I_O_2_ = 0.21) at room temperature. Mice were habituated to the plethysmography chamber and optic fiber cable for three days prior to experiments between 9:00 and 12:00 or 13:00 and 16:00. Experiments took place the next day during the same time of day as the habituation phase. On the day of the experiment, calibration of the system was performed by rapidly injecting 5 ml of air into the chamber with a syringe. Unanesthetized mice were gently handled to measure body temperature (Physitemp, Clifton, NJ, USA) and were tethered to the fiber optic cable and placed in the plethysmograph chamber. Animals were given one hour to acclimatize to the environment. Pressure changes inside the chamber were recorded with a pressure transducer, amplified (PS100W-2, EMKA Technologies, France), and digitized using PowerLab 4/26 in LabChart Version 8 (ADInstruments, Colorado Spring, CO, USA). Respiratory frequency (*f*_R_) and tidal volume (V_T_) were obtained from the plethysmograph signal. Barometric pressure, body temperature (T_b_), chamber temperature and humidity were measured to correct and standardize V_T_ and values were expressed in ml BTPS. Once the mouse appeared calm (respiratory rate ~ 150-200 breaths/min), laser stimulation (blue, wavelength = 473 nm) was performed using various settings under control conditions. Settings included 20 ms pulses repeated 5 times with a frequency of 30, 40, 50, 60 and 80 Hz and power of 10 mW to surpass the normal respiratory rate of the animal. Each train of 5 pulses were repeated for 10 seconds.

Following stimulation phases under control conditions, respiratory variables were recorded for 10 minutes as a baseline condition before mice were getting an intraperitoneal injection of either saline or the μ-opioid receptor agonist fentanyl (0.3mg/Kg). This dose of fentanyl is considered a high dose in mice (Fujii *et al*., 2019). Five minutes were then given for fentanyl to create a respiratory rate depression and laser stimulation was once again performed using various settings. Settings included 20 ms pulses repeated 5 times with a frequency of 30, 40, 50, 60 and 80 Hz and power of 10 mW. Each train of 5 pulses were repeated for 10 seconds. Mice were then anesthetized with isoflurane (3% in 50% oxygen + 50% medical air), euthanized with intracardiac injection of T-61 and brain was harvested for post-mortem histology.

Data was extracted from LabChart Version 8 (ADInstruments, Colorado Spring, CO, USA) and exported to Microsoft Excel and GraphPad Prism 9 (Version 9.3.1; GraphPad Software) for analysis. For each laser stimulation, respiratory frequency, tidal volume and inspiratory / expiratory durations were averaged over the whole stimulation period and normalized according to the preceding baseline period (frequency and tidal volume; average of 1 min) or the average of the 5 preceding breathing cycles (inspiratory and expiratory durations). These parameters were once again expressed as percentage change of the values of the preceding baseline period. The behavioural state of the animal was defined before each stimulation. The animal was considered in a calm state if respiratory rate was ≤ 125% of baseline respiratory rate and in an active state if respiratory rate was > 125% of baseline respiratory rate.

### Respiratory and behavioral profiling

Whole-body plethysmography chambers with transparent platforms were mounted on a box with an HD 1080P high-definition camera placed at the bottom of the box. Mouse movements were tracked from below for the entire duration of experiments. Videos were recorded using Pinnacle Studio 24 MultiCam Capture software (Corel, Ottawa, Ontario, Canada), resized using Pinnacle Studio 24, and exported to EthoVision XT Version 14 (Noldus, Wageningen, the Netherlands) for analysis of movement parameters. Mouse velocity (cm/s) and activity (% of pixel change) were quantified and exported to Microsoft Excel for analysis in parallel with the respiratory parameters during stimulation in freely behaving mice. Video data was aligned with plethysmography recordings using video time stamps.

### In situ hybridization

To determine the expression of *Tac1, Vglut2* and *Oprm1* and confirm virus position through *Eyfp* and *Nk1r* expression, *in-situ* hybridization was performed in C57BL/6J, *Tac1*-IRES2-Cre-D and Vglut2-ires-cre mice which received viral injection. Mice were perfused with phosphate-buffered saline (PBS) followed by formalin and the brain was harvested and placed into formalin solution overnight at room temperature. Brains were then soaked in 20% sucrose in PBS for 24 hours followed by 30% sucrose for 24 hours. Fixed brains were frozen using Tissue-Tek O.C.T. Compound (Sakura) and dry ice and stored at −80°C. Coronal sections containing the preBötC were cut at 20 μm thickness using a cryostat (Model CM3050S, Leica Biosystems, Wetzlar, Germany) and mounted on superfrost plus slides (VWR International, Radnor, PA, USA). Sections were scanned using the Axio Scan.Z1 slide scanner (ZEISS, Germany) to confirm optical fibre location. The manufacturers’ protocol was used to perform *in-situ* hybridization (RNAscope Multiplex Fluorescent Reagent v2 Assay, Advanced Cell Diagnostics, Newark, California, USA) and sections were counterstained with DAPI. Target probes used included combinations of Mm-*Tac1* (Cat No. 410351-C2) targeting *Tac1* gene mRNA, Mm-*Slc17a6* (Cat No. 319171-C2) targeting *Vglut2* gene mRNA, Mm-*Oprm1* (Cat No. 315841) targeting *Oprm1* gene mRNA, Mm-*tacr1* (Cat No. 428781) targeting NK1 receptor gene mRNA and Mm-*Eyfp* (Cat No. 551621-C3) targeting the gene coding for eYFP. Genes are indicated in italic and with the first letter in capital in this manuscript. Tissue sections 40 μm apart were scanned using the Axio Scan.Z1 slide scanner (ZEISS, Germany). As previously described, to identify sections containing the preBötC, the Mouse Brain in Stereotaxic Coordinates (3^rd^ Edition, Paxinos and Franklin) was consulted, and anatomical markers used included (1) the nucleus tractus solitarius, (2) the nucleus ambiguous, (3) the facial nucleus, (4) the hypoglossal nucleus, (5) the external cuneate nucleus as a surface landmark, and (6) the overall shape of the section. To quantify mRNA expression, 2-3 sections containing the preBötC (extending approximately 240 μm rostral-caudal) were exported from Zen (ZEISS, Germany) to Adobe Illustrator (Creative Suite 5, Adobe) where regions of interest were drawn. Images were then exported to Fiji (ImageJ) for counting. Counts were obtained for total DAPI, *Tac1* mRNA, *Vglut2* mRNA and *Oprm1* mRNA. mRNA expression was expressed either as a percentage of total DAPI cells, total *Tac1* cells, total *Vglut2* cells or total *Oprm1* cells. Images were produced using ZEN (ZEISS, Germany).

### Statistics

Statistical analysis and graphs were performed using GraphPad Prism 9 (Version 9.3.1; Graphpad Software). Figures were prepared using Adobe Illustrator (Creative Suite 5, Adobe). Data in all figures and text are reported as means ± standard error of the mean (SEM) with individual data points also displayed. Data from male and female pups were combined as preliminary statistical analyses revealed no sex specific responses to laser stimulation. Data was first tested for normality by using a Shapiro-Wilk test. Normally distributed data was analysed with ANOVAs (one-way ANOVA, two-way ANOVA or two-way repeated measures ANOVA) or a mixed effects analysis if there were missing values. Factors considered in the analyses were the effect of group (control, *Tac1, Vglut2*), laser stimulation, laser power (mW) and stimulation phase. A simple linear regression was used to determine the relationships between percentage changes in respiratory rate, velocities or baseline respiratory rates. When ANOVA results indicated that a factor (or a factorial interaction) was significant (*P* ≤ 0.05), a Tukey’s multiple comparisons test or Šidàk’s multiple comparisons test was performed for *post hoc* analysis. ANOVA results are mainly reported in the text while results from *post hoc* tests are reported in figures using symbols.

## Results

### Stimulation of *Tac1*-expressing cells in anesthetized mice

Using a Cre-lox recombination-based approach, we expressed ChETA (**Figure 1a**) in the medulla of *Tac1* cre mice. Repetition of 5 stimulations were produced containing frequencies ranging between 5, 10, 20, 30 & 40 Hz (**Figure 1b**). As no changes in respiratory rate were observed in animals which received the virus two weeks prior to experiment day (**Figure 1c**), similar stimulations were performed on animals 4 weeks following virus injections. In this group, laser stimulation increased respiratory rate compared to control group (**Figure 1b,c**; group effect: *P=0.0003*, n=25; F_(2,22)_=11.68). Absolute results for respiratory rate are provided in **Suppl. Figure 1**. Diaphragm respiratory amplitude was unaffected by laser stimulation (**Figure 1d**; group effect: *P=0.3606*, n=25; F_(2,20)_=1.074). Inspiratory duration was not influenced by stimulation (**Figure 1e**; group effect: *P=0.0934*, n=14; F_(1,12)_=3.321) but expiratory duration did decrease compared to control group (**Figure 1f**; group effect: *P=0.1158*, n=14; F_(1,12)_=2.874). Laser stimulations at 30 Hz with power of 1, 3, 5 and 10 mW showed significant differences regarding their effect on respiratory rate (**Figure 1g**; laser power effect: *P=0.0018*, n=38; F_(3,34)_=6.201).

**Figure 1.**
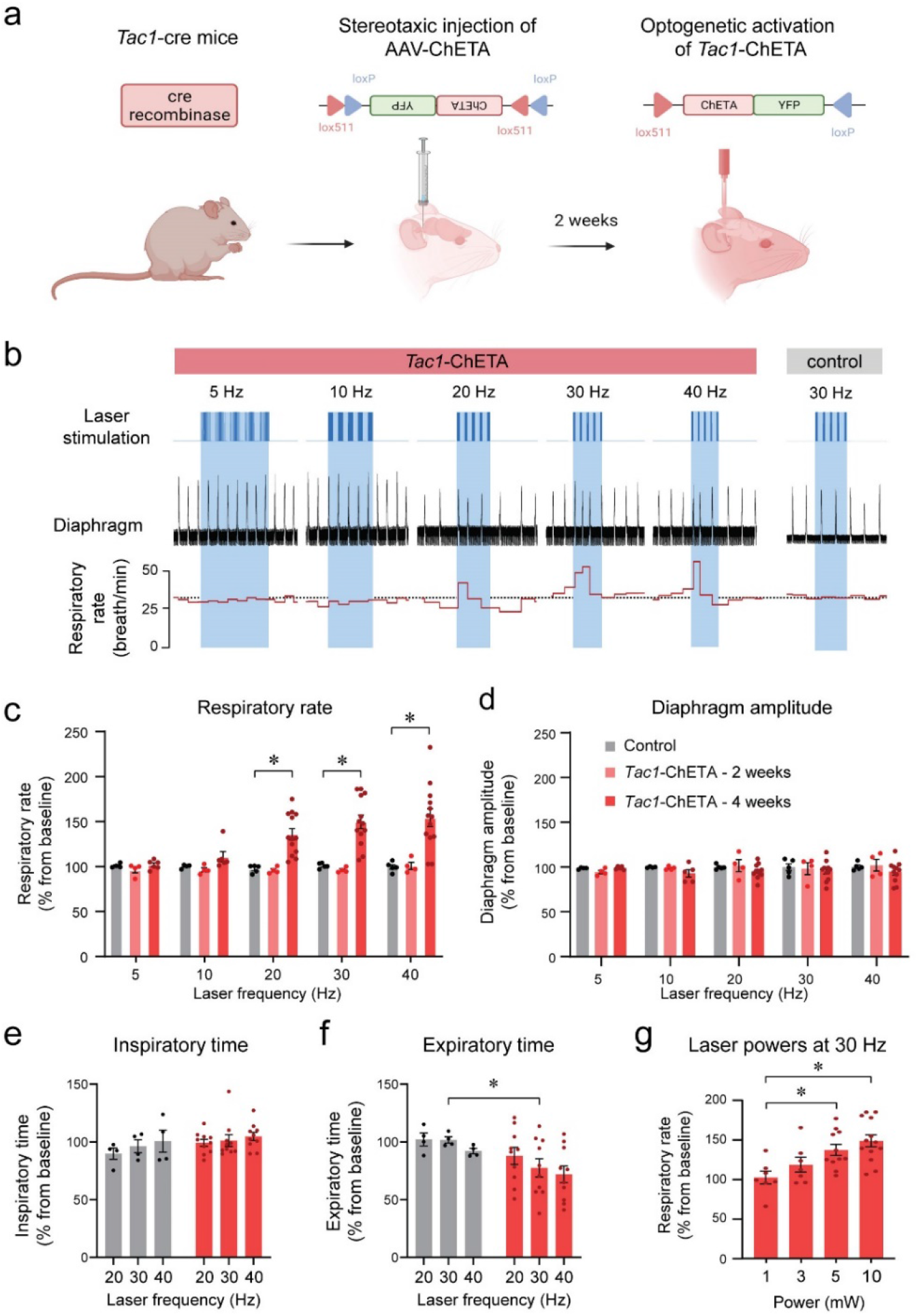
Photostimulation of *Tac1* preBötC cells increases respiratory rate in anesthetized mice. **(a)** ChETA was expressed in *Tac1* preBötC cells by injecting the AAV-ChETA^fl/fl^ virus in *Tac1* cre-expressing mice. **(b)** After two or four weeks of incubation, laser stimulations were performed at various frequencies in control and *Tac1*-ChETA anesthetized mice. **(c)** Laser stimulation increased respiratory rate at 20, 30 and 40 Hz. *Tac1* cells were only stimulated after virus incubated for 4 weeks, but not 2 weeks (n=25). **(d)** No effect was observed on diaphragm amplitude (n=25). **(e)** Laser stimulation had no effect on inspiratory time, but **(f)** significantly decreased expiratory time at 30 Hz (n=14). **(g)** Laser powers stimulated respiratory rate at 5 and 10 mW (n=38). Data are presented as means ± SEM, with individual data points. * indicate means significantly different from corresponding controls or laser powers with P<0.05. Panel A was created using Biorender.com.

### Response of *Tac1*-expressing cells depends on stimulation phase

We produced various stimulations at different time points during the respiratory cycle with high temporal specificity using optogenetics. Frequency of 30 Hz and laser power of 10 mW were used. Stimulation phase was defined as the time between the beginning of inspiration (beginning of the respiratory cycle) and the onset of the laser (**Figure 2a**). When stimulation phase is normalized to the preceding unstimulated respiratory cycle (T_b_), simulation phase represents the percentage of the respiratory cycle when the laser was turned on, with the start of inspiration defined as 0% of the cycle and the end of expiration defined as 100% of the cycle (**Figure 2b**). All induced periods (T; duration of the induced respiratory cycle) were also normalized to the preceding unstimulated respiratory cycle (T_b_) and presented as percentage of Tb (**Figure 2b**). When individual data points were plotted for each normalized stimulation phase from all animals, the induced period decreased when laser stimulation occurred past 20% of the respiratory cycle (**Figure 2b**).

**Figure 2.**
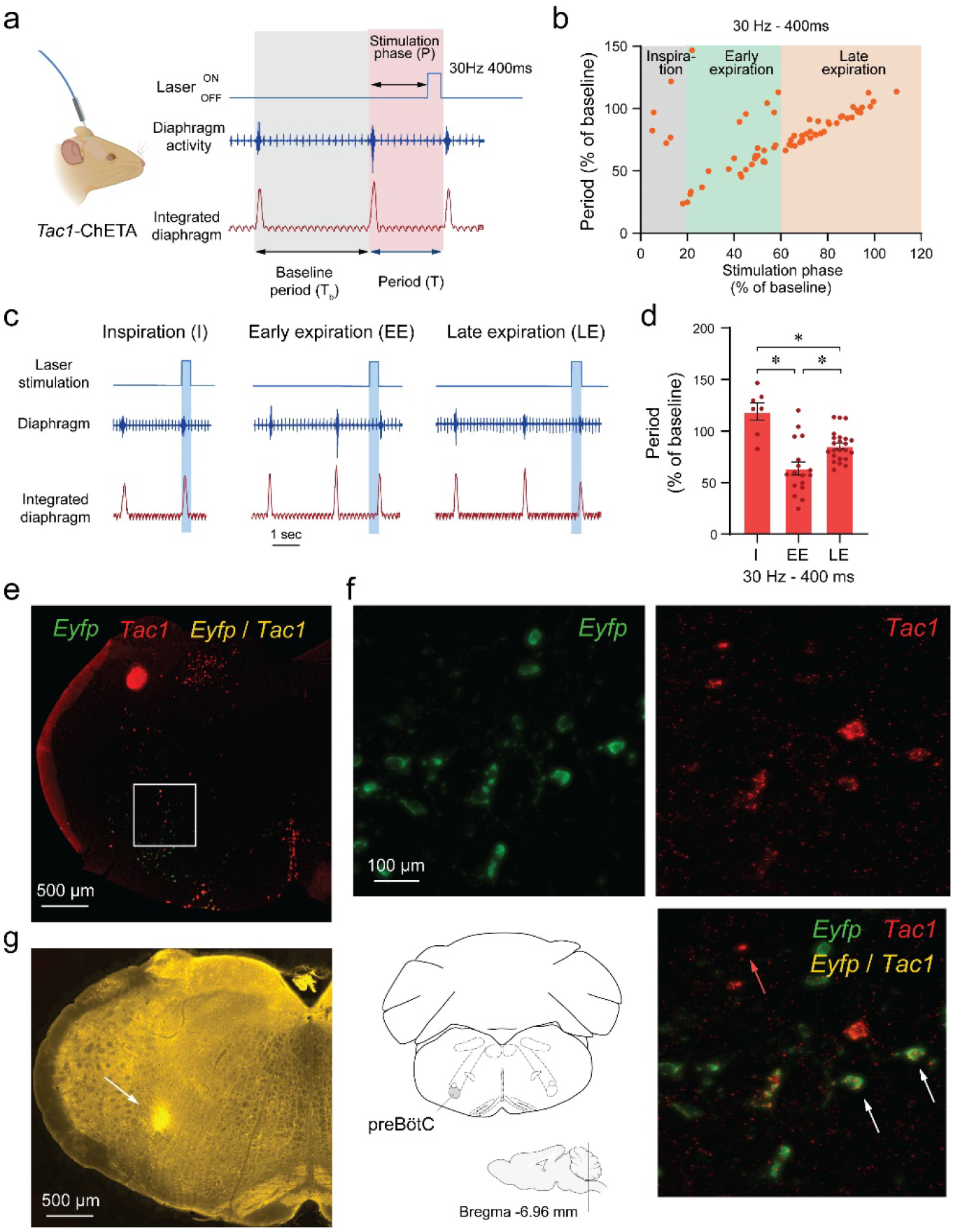
Phase-dependent photostimulation of *Tac1* preBötC cells. **(a)** For each laser stimulation, the stimulation phase (i.e. the time between the beginning of inspiration and the onset of the laser), the induced period (T), and the baseline period (T_b_) were measured. All stimulation phases and induced periods were normalized to the preceding unstimulated respiratory cycle (T_b_) and presented as percentage of Tb **(b)** The changes in periods were represented against the stimulation phase at 30 Hz. When stimulation of *Tac1* cells was performed during inspiration, period was not changed. The respiratory period was substantially reduced when stimulation occurred early in expiration and to a lesser degree late during expiration. **(c)** Laser stimulations were categorized according to their occurrences during inspiration (I), early (EE) and late expiration (LE). **(d)** *Tac1* stimulations occurring during early and late expirations significantly reduced the respiratory period (n=47). **(e)** Viral injection in the preBötC and opsin channel ChETA expression in *Tac1* cells were confirmed using *in-situ* hybridization. **(f)** *Tac1* (red) and *Eyfp* (green) mRNA were found in the preBötC area with co-expression of both mRNAs (yellow). **(g)** The locations of the optical fiber placement above the preBötC were confirmed with post-mortem histology. Data are presented as means ± SEM, with individual data points. * indicate means significantly different from stimulation phases with P<0.05. Panel A was created using Biorender.com.

Optical stimulation phases were then grouped under three categories according to their timing in the respiratory cycle: inspiration, early expiration and late expiration (**Figure 2c**). Stimulation occurring during inspiration had no effects on the period (118% of baseline period; **Figure 2b, d**). However, stimulation decreased the period when it occurred early in expiration phase (63% of baseline period) and also decreased the period when it occurred late in expiration phase, showing an immediate response in this phase of the respiratory cycle (85 % of baseline period) (**Figure 2b, d**; stimulation phase effect: *P<0.0001*, n=47; F_(2,44)_=18.37). Post-mortem histology confirmed that the optical fibre was positioned in regions of the medulla, dorsal to the preBötC allowing the laser light to penetrate ventral to the optical fibre (**Figure 2g**). We confirmed viral injection in the preBötC and opsin channel ChETA expression in *Tac1* cells through *in situ* hybridization with probes targeting *Tac1* (red) and *Eyfp* (green). All cells tagged with *Eyfp* (expressing ChETA) had substantial expression of *Tac1* mRNA (**Figure 2e, f**). Importantly, ChETA was expressed in a region of the medulla rich in NK-1Rs consistent with the preBötC (**Supp. Figure 2**). In conclusion, photostimulation of *Tac1* preBötC triggered inspiration when occurring during expiration, but not during inspiration.

### *Tac1*-expressing cells: a glutamatergic subpopulation in the preBötzinger complex

Cells expressing *Tac1* have previously been localized in the preBötzinger region using *in-situ* hybridization (Sun *et al*., 2019). In this region, they show a strong co-expression with NK-1R-expressing cells, a gene coding for neurokinin-1 receptors found in the preBötC and known to stimulate breathing once activated by substance P (Gray *et al*., 1999). Knowing the *Tac1*-expressing cells targeted in this study are mainly excitatory as demonstrated in **Figure 1**, we aimed to determine whether they can be considered as a subpopulation of glutamatergic cells. Using *in-situ* hybridization, we found presence of *Tac1*-expressing cells (red) in both the NTS and preBötC as well as the medullary raphe (**Figure 3a-d**). In the preBötC region, *Tac1* cells overlapped were expressed in 11% of DAPI cells, while Vglut2 cells in 44% of DAPI cells (**Figure 3e**). Cells co-expressing *Vglut2* (green) and *Tac1* (red) were found in 11% of DAPI cells. *Tac1* cells co-expressing *Vglut2* constituted 93% of *Tac1* cells, but only 24% of *Vglut2* cells (**Figure 3f**). Importantly, a vast majority of *Tac1* neurons are found in the preBotC with little expression in adjacent motor nuclei (**Suppl. Figure 3**). These results suggest that a large majority of *Tac1* cells are excitatory glutamatergic preBötC neurons.

**Figure 3.**
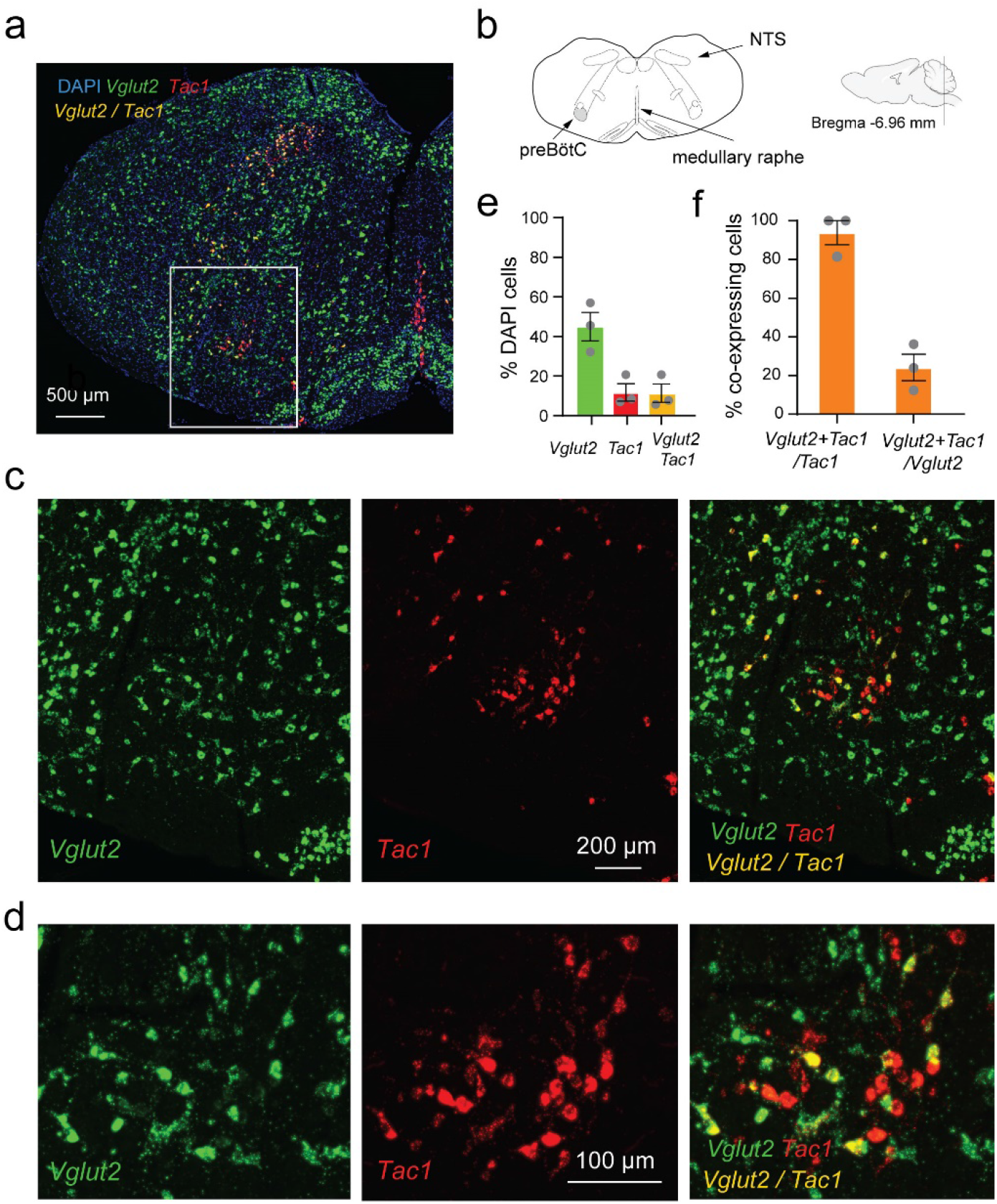
*Vglut2* and *Tac1* mRNA expressions in the medulla. **(a)** *Vglut2* (green) and *Tac1* (red) mRNAs are expressed in the region of the preBötC as shown by *in-situ* hybridization in wild type mice. **(b)** Substantial expression of *Vglut2* mRNA was observed in the medulla with a cluster of *Tac1* in the preBötzinger Complex (preBötC). **(c)** In the preBötC, part of the cells expressing *Tac1* also expressed *Vglut2* (co-expression shown by yellow). **(d)** Magnified views of the central part of panel c. **(e)** In the preBötC, about 44% of the cells (shown with DAPI) expressed *Vglut2*, 11% *Tac1* alone, and 11% co-expressed *Tac1* and *Vglut2*. A total of 3 mice were used. **(f)** The majority of *Tac1* cells co-expressed Vglut2 (93%), whereas only 24% of Vglut2 cells co-expressed Tac1. DAPI was shown in blue.

### Stimulation of *Vglut2*-expressing cells in anesthetized mice

Knowing that *Tac1*-expressing cells also co-express *Vglut2*, we characterized the role of *Vglut2* preBötC neurons in breathing. Using a similar approach as above, we expressed the excitatory ChETA in *Vglut2* cells (**Figure 4a**). We produced repetitions of 5 stimulations with various frequencies (5, 10, 20, 30 & 40 Hz) (**Figure 4b**). Stimulations increased respiratory rate compared to control group (**Figure 4b, c**; group effect: *P=0.0019*, n=11; F_(1,9)_=18.77). Absolute results for respiratory rate are provided in **Suppl. Figure 4**. Diaphragm respiratory amplitude was unaffected by laser stimulation (**Figure 4d**; group effect: *P=0.5527*, n=11; F_(1,9)_=0.3803). We then determined whether inspiratory or expiratory durations were affected by laser stimulation. Inspiratory duration was not influenced by stimulation (**Figure 4e**; group effect: *P=0.1476*, n=11; F_(1,8)_=2.569). Therefore, the increased respiratory rate can be mainly explained by the decrease in expiratory duration compared to control group (**Figure 4f**; group effect: *P=0.0057*, n=11; F_(1,8)_=14.03), as observed with stimulation of *Tac1* neurons. Different power of laser stimulations (1, 3, 5 and 10 mW) showed significant differences on respiratory rate at 40 Hz (**Figure 4g**; power effect: *P=0.0314*, n=24; F_(3,20)_=3.605). To confirm viral injection in the preBötC and ChETA expression in *Vglut2* cells, we performed *in situ* hybridization with probes targeting *Vglut2* (violet) and *Eyfp* mRNA (green). All cells tagged with *Eyfp* (shown in green) were also co-expressing *Vglut2* (in purple) as shown by white cells (**Figure 4h**) in the region of the preBötC, confirming accurate expression of opsin channels in the preBötC.

**Figure 4.**
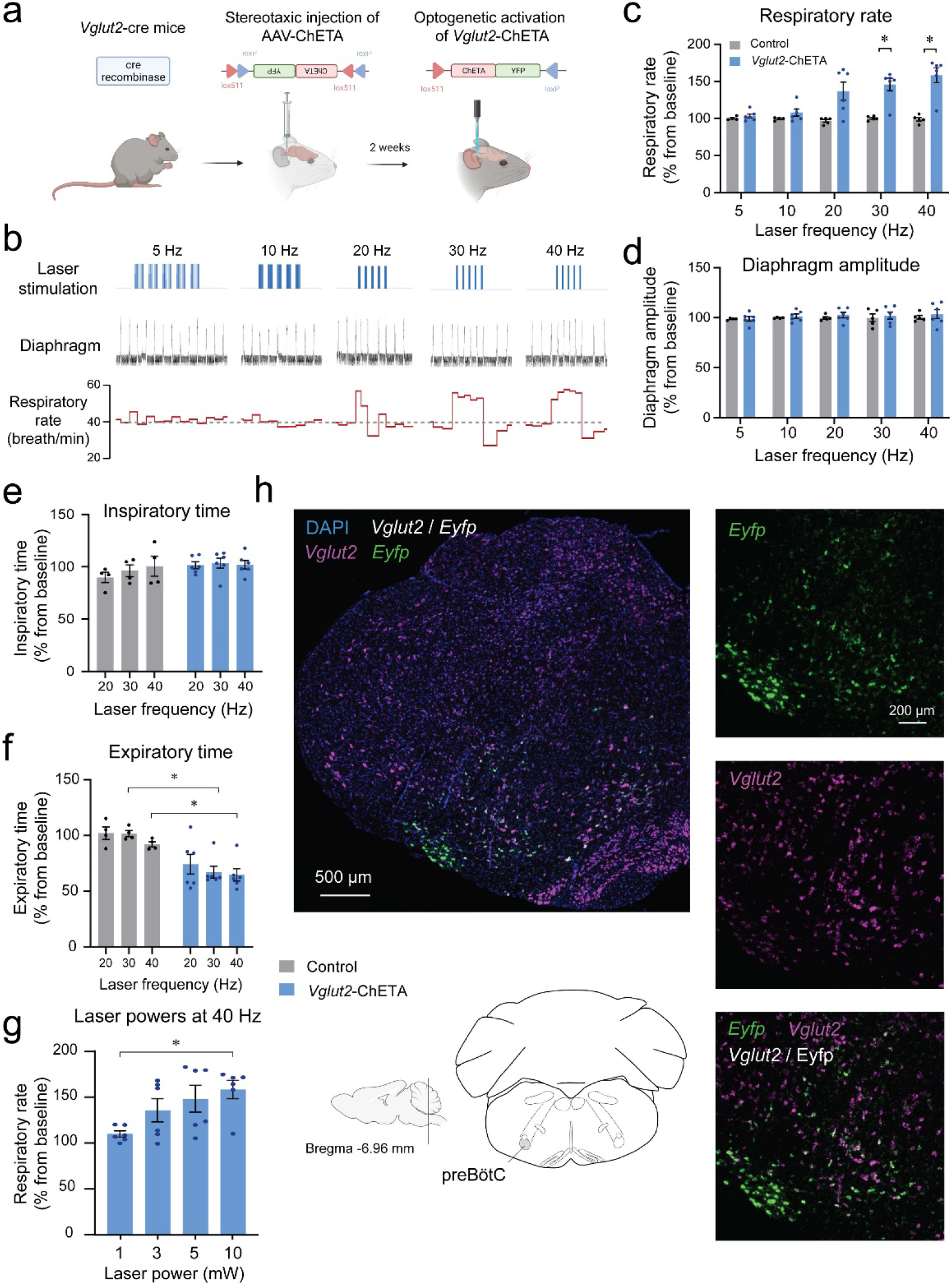
Photostimulation of *Vglut2* preBötzinger Complex cells increases respiratory rate in anesthetized mice. **(a)** The adeno-associated virus AAV-ChETA^fl/fl^ was injected into the preBötC of *Vglut2* cre-expressing mice. After two weeks of incubation with AAV-ChETA, *Vglut2* cells expressed ChETA and YFP. **(b)** In anesthetized mice, diaphragm muscle activity was recorded and laser stimulations at various frequencies were performed in the preBötC. **(c)** Respiratory rate was significantly increased by laser stimulation at 30 and 40 Hz and **(d)** no effect was observed to diaphragm amplitude (n=11). Increased respiratory rate was not due to decrease in **(e)** inspiratory time, but rather decreased **(f)** expiratory time (n=11). **(g)** Respiratory rate was increased by laser stimulation at 10 mW (n=24). **(h)** *In situ* hybridization was performed in animals injected with AAV-ChETA and showed that ChETA (marked with *Eyfp* mRNA) was expressed in the region of the preBötC and co-expressed with *Vglut2*. Data are presented as means ± SEM, with individual data points. * indicate means significantly different from corresponding controls or laser power with P<0.05. Panel A was created using Biorender.com.

### Stimulation of *Tac1* preBötC cells in freely behaving mice

To determine the role of *Tac1* preBötC cells in promoting breathing *in vivo*, photostimulation of *Tac1* preBötC cells was performed in freely-behaving rodents while recording respiratory activity (**Figure 5a**). Based on baseline respiratory rate, animal state when laser stimulation happens were divided between calm and active state (**Figure 5b)**. *Tac1*-expressing preBötC cells were stimulated using a range of frequencies and respiratory rate was increased at 30, 40, 50, and 60Hz, but not at 80 Hz (**Figure 5c**; group effect: *P=0.0037*, n=14; F_(1,12)_=12.90) by a combined decrease in both inspiratory time (**Figure 5c**; group effect: *P=0.0002*, n=14; F_(1,12)_=26.82) and expiratory time (**Figure 5c**; group effect: *P=0.0156*, n=14; F_(1,12)_=7.920). Absolute results for respiratory rate are provided in Supplementary Figures. When photostimulation occurred when the animal was in calm state (as defined by low baseline respiratory rate), photostimulation significantly increased respiratory rate compared to controls while only moderate effect was observed in active states (**Figure 5d**; group effect: *P<0.0001*, n=49; F_(3,46)_=14.97). This increase in respiratory rate was negatively correlated with the baseline respiratory rate (**Figure 5e**; R^2^=0.1802, *P=0.0273*, n=27; F_(1,25)_=5.494). To better account for the differences in baseline respiratory rate observed between calm and active states, we normalized respiratory rate according to respiratory rate preceding photostimulation. During calm states, photostimulation increased respiratory rate, whereas no effect was observed during active state (**Figure 5f**; group effect: *P=0.0014*, n=30; F_(1,5)_=40.79). In summary, our results demonstrated that stimulation of *Tac1* preBötC cells promote substantially respiratory rate during calm state, but only moderately during active state.

**Figure 5.**
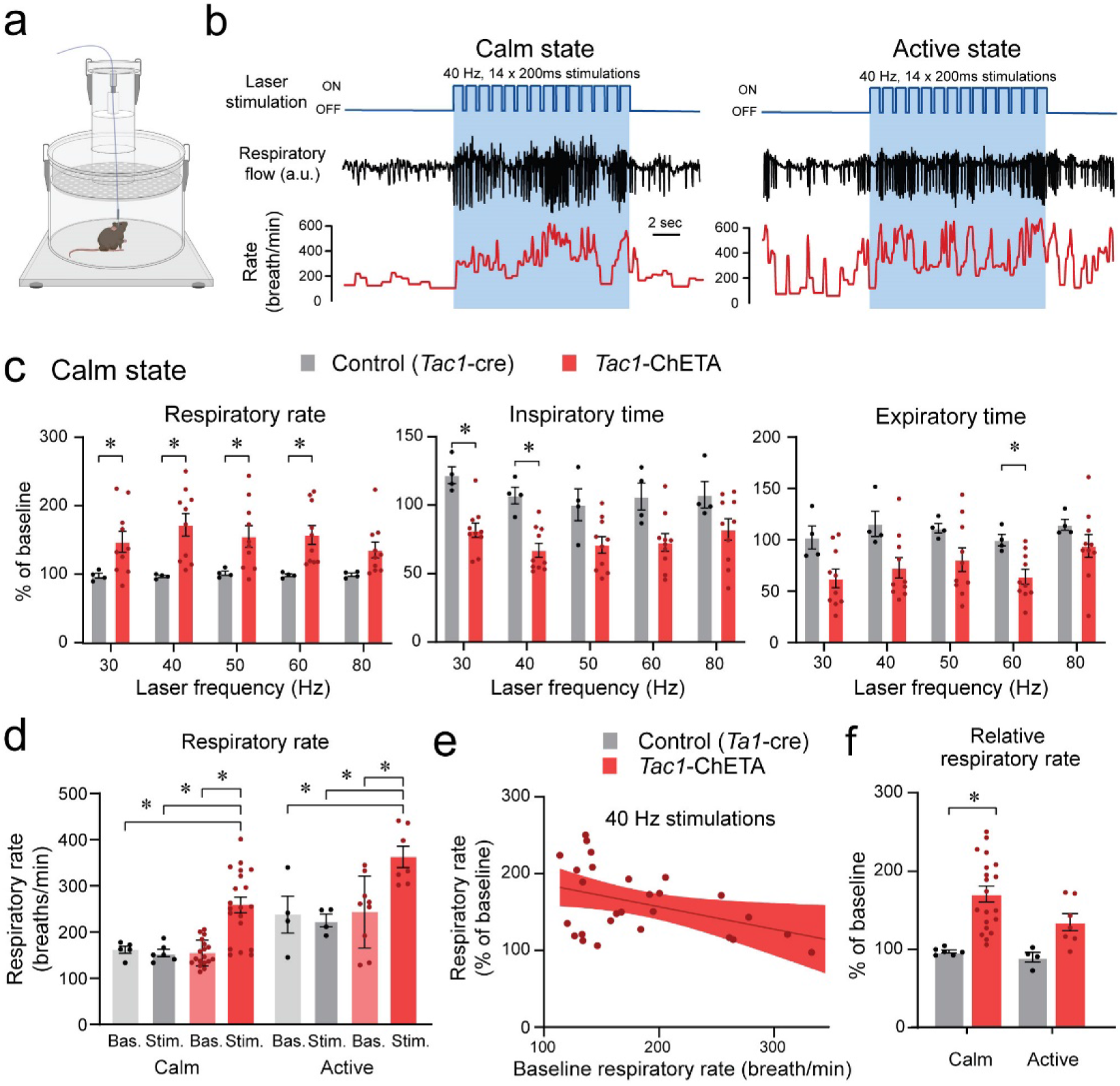
State-dependent respiratory changes by photostimulation of *Tac1* preBötC cells in freely-behaving mice. **(a)** Using a similar approach than above, ChETA was expressed in *Tac1* preBötC cells and respiratory activity was measured using whole-body plethysmography. **(b)** Representative tracings of laser stimulations at 40Hz where stimulations happen with the animal considered in a calm or active state. The effects of photostimulation were reversed when stimulations were stopped. **(c)** In calm state, *Tac1* cell stimulations at 30, 40, 50, and 60 Hz increased rate due to a combination of decreased inspiratory and expiratory times (n=14). **(d)** Stimulation of *Tac1* neurons strongly increased absolute respiratory rate in calm state whereas in active states it had a moderate effect. **(e)** Increased respiratory rate was negatively correlated with the baseline respiratory rate. **(f)** To determine the state-dependent effects of photostimulation, respiratory rate changes were expressed as a function of the baseline respiratory rate (before laser stimulation). When baseline respiratory rate was relatively low, photostimulation substantially increased respiratory rate, whereas photostimulation was not as effective when baseline respiratory rate was high (n=27). Data are presented as means ± SEM, with individual data points. * indicate means significantly different from corresponding controls with P<0.05. Panel A was created using Biorender.com.

### Motor hyperactivity induced by stimulation of *Tac1*-expressing cells

To determine whether stimulation of *Tac1*-expressing cells induces behavioral changes, we assessed locomotor activity using a video recording system and tracking software (**Figure 6a**). Representative heat map of both control mice and mice with photostimulation of *Tac1* cells are shown in **Figure 6b** during baseline, laser stimulation and recovery phases. Mice quickly responded to a 10 sec stimulation period at 40Hz with an increase in movements (**Figure 6c**). Mouse activity in response to stimulation (percentage of pixel change in the area recorded) was significantly higher than control group through both stimulation and recovery phases (**Figure 6d**; group effect: *P=0.0476*, n=14; F_(1,12)_=4.869). Velocity was similarly increased by photostimulation compared to control animals (**Figure 6e**, group effect: *P=0.0122*, n=14; F_(1,12)_=8.687). While respiratory recordings by whole-body plethysmography is the best approach to assess respiratory activity in freely-moving and non-anesthetized rodents, our system also captures behaviors that may impact respiratory recordings (Montandon & Horner, 2019). To better assess the relationships between locomotor behaviors and breathing, we correlated respiratory rate and velocity in freely behaving mice in response to *Tac1* photostimulation. During *Tac1* photostimulation, velocity was positively correlated with respiratory rate (**Figure 6f**; R^2^=0.5623, *P=0.0125*, n=10; F_(1,8)_=10.28), suggesting that higher respiratory rate was associated with higher locomotor activity. In control animals, increased respiratory rate was not associated with increased velocity, suggesting that an association between velocity and respiratory rate was only apparent when *Tac1* preBötC cells were stimulated.

**Figure 6.**
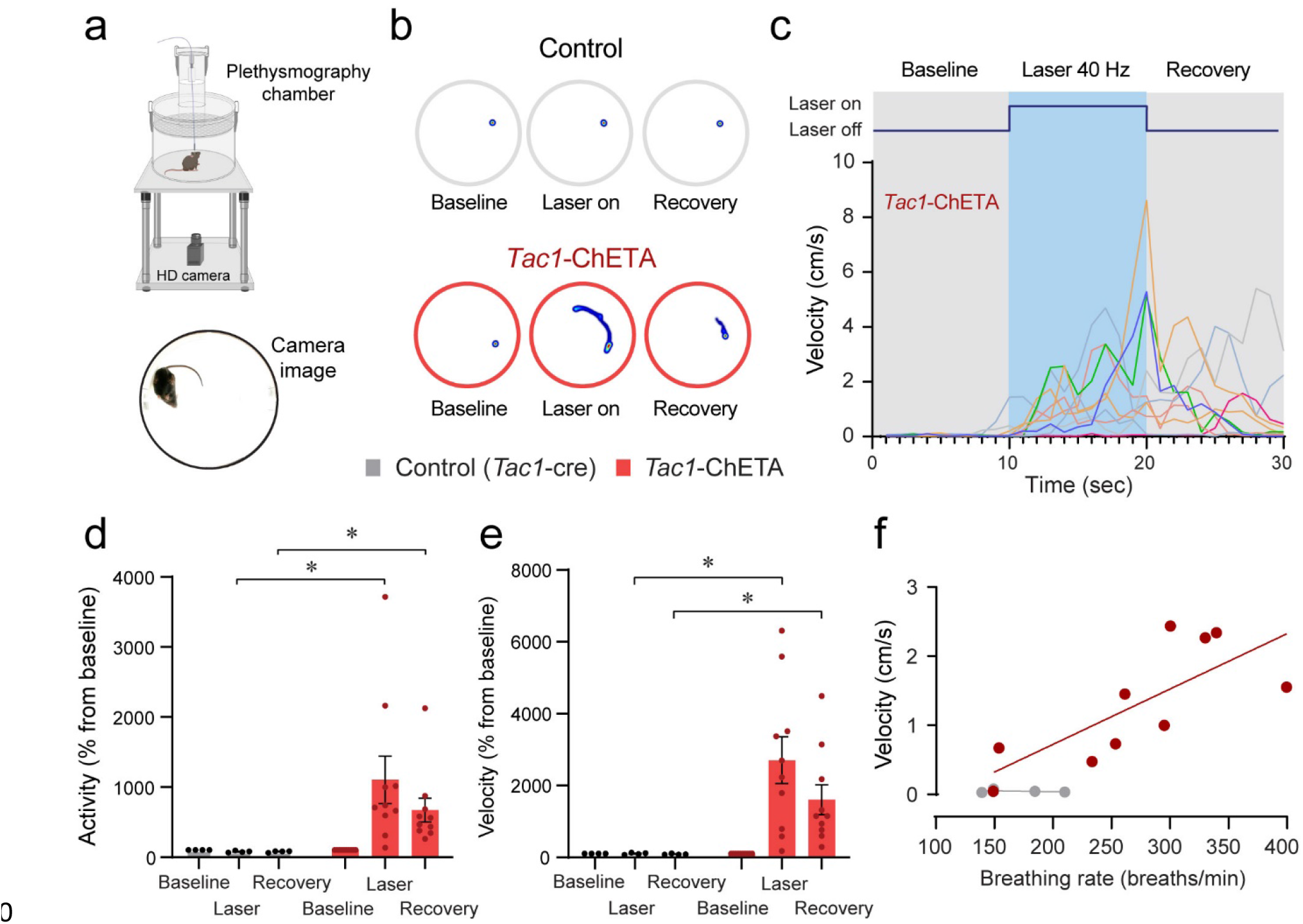
Optogenetic stimulation of *Tac1* preBötC cells promotes locomotion in freely-behaving mice. **(a)** Locomotor activity in freely behaving *Tac1-*cre mice expressing ChETA was assessed with a high-definition camera. **(b)** Heat maps showing the locomotion of control and *Tac1*-ChETA mice in 3 different conditions; before stimulation (baseline), during stimulation (laser ON), and following stimulation (recovery). Each circle represents 10-sec locomotion under the corresponding condition. **(c)** The velocity (cm/sec) for each *Tac1*-ChETA animal strongly increases during stimulation at 40Hz followed by a reduction in velocity with cessation of stimulation. **(d)** Activity (pixel changes inside the recording circle) also increased significantly with laser stimulation in *Tac1*-ChETA but not in control mice (n=14). **(e)** This effect was mainly due to substantial increases in velocity in *Tac1*-ChETA compared to control mice. This effect was sustained for a few seconds during recovery (n=14). **(f)** Correlations between velocity and respiratory rate showed that increases in respiratory rate due to *Tac1* stimulation were associated with increased velocities (n=10). Data are presented as means ± SEM, with individual data points. * indicate means significantly different from corresponding controls with P<0.05. Panel A was created using Biorender.com.

### Co-expression of *Tac1* and *Oprm1* in the preBötzinger complex

Considering the role of *Tac1*-expressing cells in rhythmic breathing and knowing that the preBötC plays a major part in respiratory depression by opioid drugs (Montandon *et al*., 2011; Stucke *et al*., 2015), we determined whether *Tac1* mRNA is co-expressed with *Oprm1* (gene for MORs) mRNA in the preBötC. We performed *in situ* hybridization in medullary sections containing the preBötC in wild type mice (C57BL/6J) (**Figure 7a, b**). In the region of the preBötC, *Oprm1* was expressed in 40.5 ± 6.8% of DAPI-stained cells while *Tac1* was expressed in 21.8 ± 8.9% of DAPI-stained cells (**Figure 7c**; n=3). Co-expression of both *Oprm1* and *Tac1* was found in 17.7 ± 7.6% of DAPI-stained cells (**Figure 7c**; n=3). Interestingly, most cells expressing *Tac1* also co-expressed *Oprm1 (*75.9 ± 5.7% of *Tac1*-expressing cells; **Figure 7c, d**; n=3) but only a small fraction of *Oprm1* cells were expressing *Tac1* (38.3 ± 13.7% of *Oprm1-*expressing cells; **Figure 7c, d**; n=3). In summary, *Tac1* cells constitute a small fraction of cells found in the preBötC but a large proportion of these cells co-express *Oprm1* mRNA.

**Figure 7.**
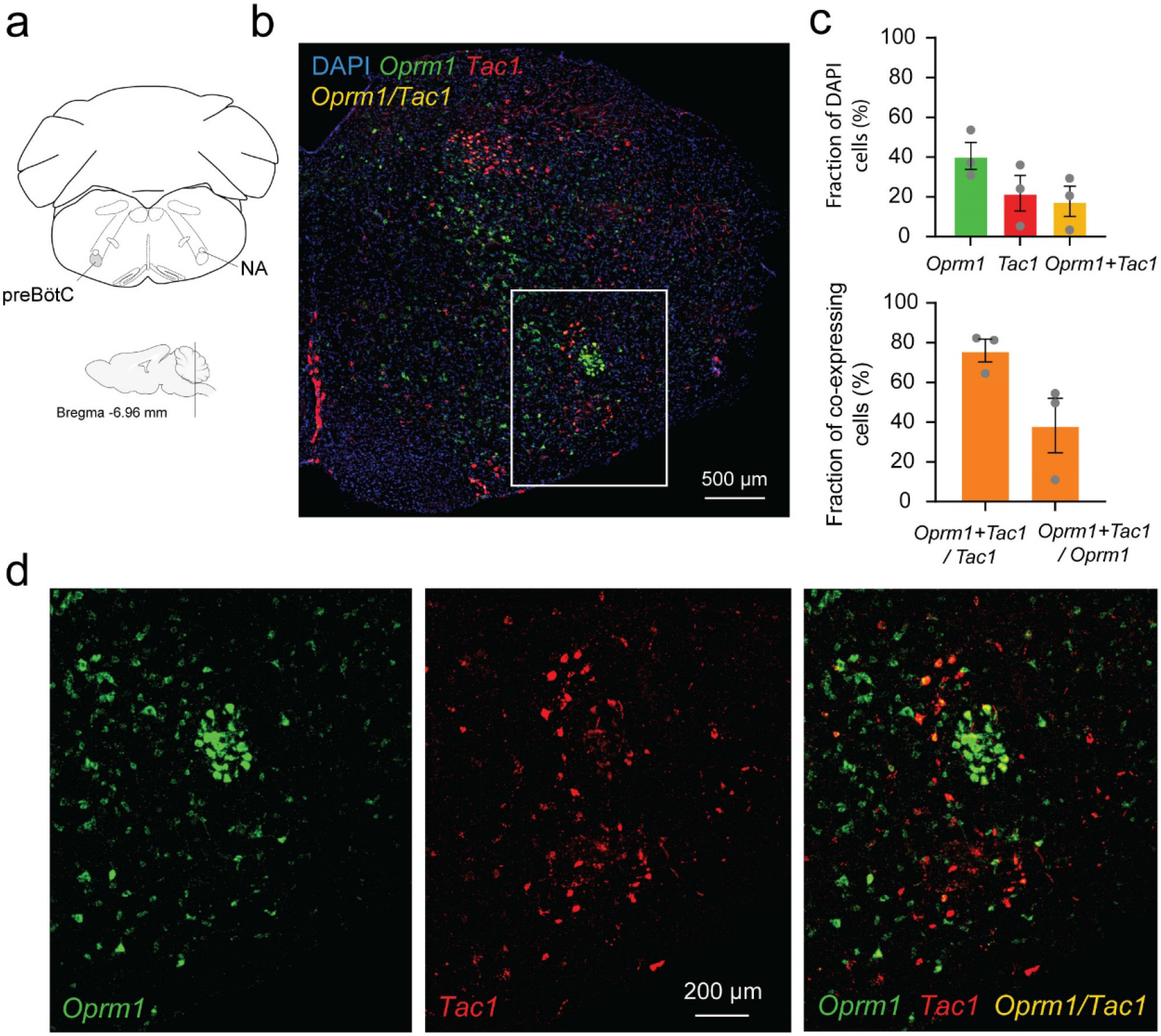
Co-expression of *Oprm1* and *Tac1* mRNAs in preBötC cells. **(a)**. *In-situ* hybridization was performed on sections containing the preBötC about 6.96 mm caudal to Bregma. **(b)** In the medulla, *Oprm1* (the gene encoding for MOR shown in green), *Tac1* (the gene for Substance P in red) and DAPI (in blue) were widely expressed in the preBötC. **(c)** Cell counting in the preBötC shows that about 40.5% of the cells contained *Oprm1* and 21.8% contained *Tac1* (n=3). In the preBötC, 75.9% of cells expressing *Tac1* also co-expressed *Oprm1* (n=3). Conversely, about 38% of cells expressing *Oprm1* also co-expressed *Tac1* (n=3). **(d)** In a magnified view of the preBötC, *Oprm1* mRNAs formed a cell cluster in the ventral part of the medulla and co-expressed *Tac1*. NA, nucleus ambiguous.

### Stimulation of *Tac1*-expressing cells reverses opioid-induced respiratory depression

Considering that photostimulation of *Tac1* cells increased breathing and that these cells co-expressed *Oprm1*, we aimed to determine whether stimulation of *Tac1*-expressing cells can reverse opioid-induced respiratory depression. We injected the opioid fentanyl (intramuscular; 5μg/kg) in anesthetized mice followed by stimulations of 20, 30 and 40 Hz (**Figure 8a, b**). As expected, fentanyl injection reduced respiratory rate to 53.6% of baseline value and laser stimulation with frequencies of 20, 30 and 40Hz significantly reversed fentanyl-induced respiratory rate depression (**Figure 8c, d**; stimulation effect: *P=0.0002*, n=35; F_(4,30)_=7.986, **Suppl. Figure 5**). To determine the effect of photostimulation on respiratory depression by opioid drugs, in more realistic conditions and without anesthetics, we performed similar experiments in freely-behaving rodents. We used whole-body plethysmography to assess respiratory responses to the opioid fentanyl (0.3mg/kg, intraperitoneal) (**Figure 8e**). To take into account the stress associated with intraperitoneal injection in live mice, saline injections were used as controls. In mice injected with saline, respiratory rate increased following injection due to mouse handling and stress and photostimulation at 40 Hz did not significantly change respiratory rate. Injection of fentanyl did not increase respiratory rate, and showed a significant lower respiratory rate compared to mice injected with saline. Photostimulation significantly increased respiratory rate (**Figure 8f, g**; group x stimulation effect: *P<0.0001*, n=14; F_(2,24)_=16.84), despite fentanyl lowering respiratory rate compared to saline. Interestingly, saline injection did not change tidal volume while fentanyl injection increased it. Photostimulation did not change tidal volume (**Figure 8f, g**; group x stimulation effect: *P<0.0001*, n=14; F_(2,24)_=16.42). Mouse movements were then analysed following saline or fentanyl injection. Fentanyl injections increased velocity in mice and laser stimulation further raised it (**Figure 8h**; group x stimulation effect: *P=0.0941*, n=13; F_(2,22)_=2.637). In conclusion, fentanyl presented a lower respiratory rate compared to saline injection, which was fully reversed by stimulation of *Tac1* preBötC neurons in anesthetized and freely-behaving mice.

**Figure 8.**
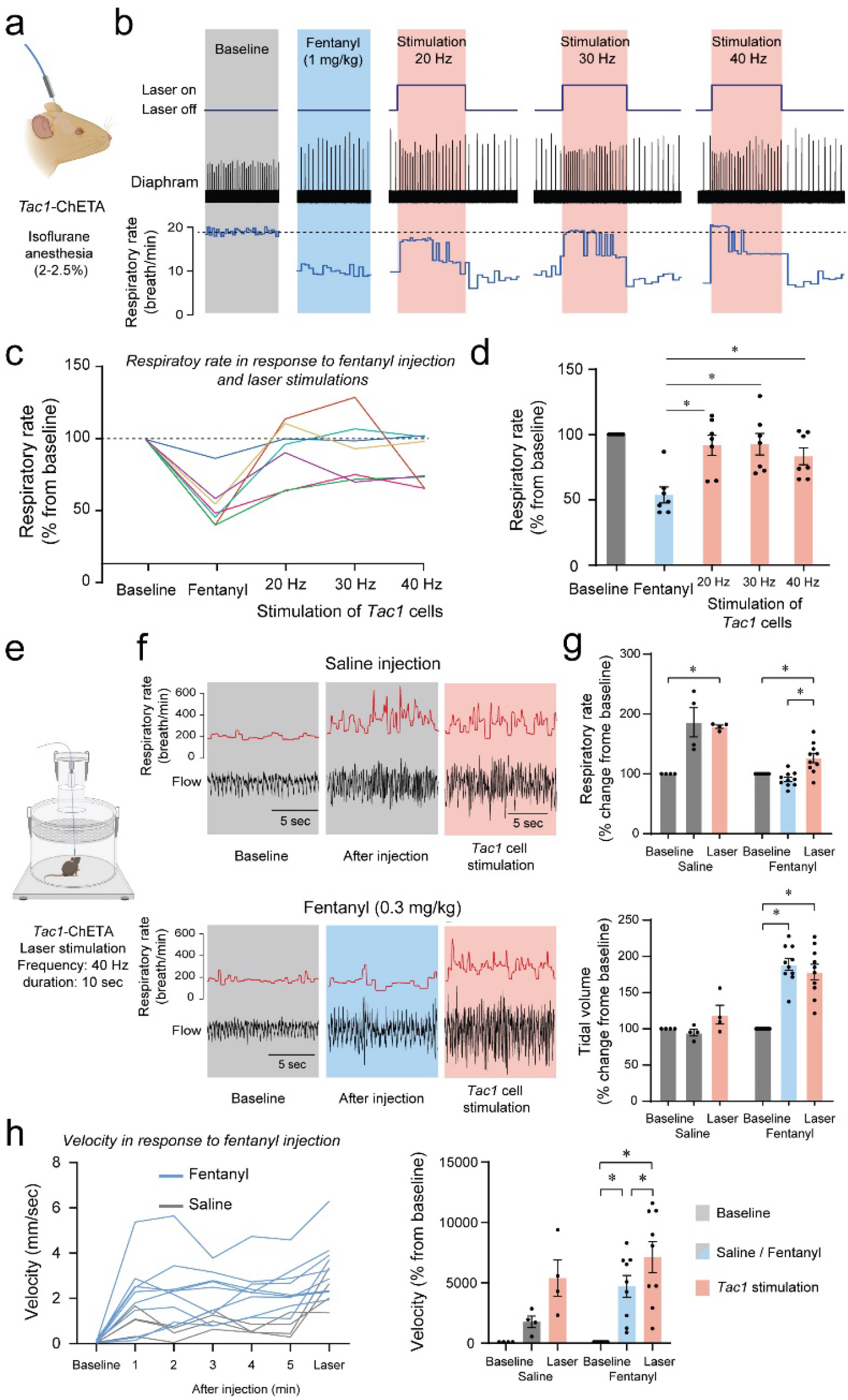
Photostimulation of *Tac1* preBötC cells reverses respiratory depression by the opioid fentanyl. **(a)** Injection of fentanyl (5µg/kg) performed in anesthetized *Tac1*-ChETA mice. **(b)** Injection of fentanyl depressed breathing, with this effect reversed by photostimulation of *Tac1* preBötC cells at 20, 30 and 40 Hz. **(c)** All 7 animals receiving fentanyl showed respiratory rate depression by fentanyl reversed by Tac1 photostimulation (each color represents a separate tracing for each animal). Once the laser was turned off, breathing rate returned to low breathing rates due to fentanyl. **(d)** Mean data showed that stimulation of *Tac1* cells reversed respiratory depression at each laser frequency used (20, 30, 40 Hz, n=28). **(e)** In freely-moving, non-anesthetized, *Tac1-*ChETA mice, respiratory responses to the opioid fentanyl (0.3mg/kg, intraperitoneal) was assessed using whole-body plethysmography. **(f)** Representative tracings of diaphragm activity and respiratory rate showed that fentanyl depressed respiratory rate compared to saline and that Tac1 stimulation reversed respiratory depression. **(g)** Mean data show how fentanyl significantly reduced respiratory rate observed with saline injection, an effect reversed by Tac1 stimulation in fentanyl conditions only (n=14). Tidal volume was increased by fentanyl injection but unaffected by photostimulation. **(h)** Locomotor activity was assessed and showed increase in velocity following injection and stimulation at 40 Hz (n=13). All absolute values (not normalized according to baseline) of respiratory rate and tidal volume can be found in **Suppl. Figure 4** and are consistent with normalized results. Data are presented as means ± SEM, with individual data points. * indicate means significantly different from corresponding controls with P<0.05. Panel A, E were created using Biorender.com.

## Discussion

Characterizing the neuronal elements of the preBötzinger Complex is key to understand the complexity of the network generating breathing and to identify therapeutic targets when breathing fails. We characterized a subpopulation of glutamatergic neurons expressing the *Tac1* gene that promotes breathing and can be targeted to reverse respiratory depression by opioid drugs. By unilaterally modulating *Tac1* preBötC cells, we modulated rhythmic breathing in a spatially and temporally precise manner in both anesthetized and freely moving mice. This is the first *in vivo* evidence that optogenetic activation of *Tac1* preBötC cells increases breathing, resets the inspiratory cycle in a phase dependent manner, and induces a strong behavioral response. *Tac1* role on breathing depends on the state of the animal (calm *vs*. active), and therefore the overall excitability of the respiratory network, as a minimal effect of *Tac1* stimulation was observed when the animal was behaviorally active.

### *Tac1*-expressing cells, a small subpopulation of glutamatergic cells, regulate breathing

The preBötzinger complex is a heterogeneous network containing multiple subpopulations of molecularly defined neurons balancing excitation and inhibition to produce rhythmic breathing. In the preBötC, most inspiratory-related excitatory cells are glutamatergic and their activation is essential to initiate synchronized rhythmic activity and produce a breath (Wallen-Mackenzie *et al*., 2006). Stimulation of glutamatergic excitatory cells in the preBötC entrains respiratory activity both *in vitro* and *in vivo* (Baertsch *et al*., 2018; Oliveira *et al*., 2021). However, much remains unknown about the different subclasses of glutamatergic preBötC neurons and how they produce rhythmic breathing. A large subset of excitatory preBötC neurons expressed the transcription factor developing brain homeobox 1 protein (Dbx1) and are potential drivers of the preBötC rhythm (Baertsch *et al*., 2018; Vann *et al*., 2018). In the present study, we focused on a limited group of excitatory neurons expressing the neuropeptide substance P (encoded by the *Tac1* gene). Substance P is highly abundant and widely spread in the central and peripheral nervous system including the brainstem and the preBötC. It is involved in respiratory, cardiovascular and gastrointestinal regulation as well as nociception and chemoreception but the function of Substance P-expressing cells in the regulation of breathing *in vivo* is unknown (Otsuka & Yoshioka, 1993; Gray *et al*., 1999; Hokfelt *et al*., 2001; Pena & Ramirez, 2004). *In vitro*, substance P increases respiratory rhythm by activating neurokinin-1 receptors or NK-1R (Gray *et al*., 1999; Liu *et al*., 2004; Pena & Ramirez, 2004; Yeh *et al*., 2017; Sun *et al*., 2019). NK-1Rs are G-protein-coupled receptors stimulating rhythmic breathing through GIRK channels (Montandon *et al*., 2016a). NK-1R-expressing preBötC cells are predominantly glutamatergic and mediates inspiratory activity (Guyenet *et al*., 2002). Interestingly, destruction of NK-1R-expressing preBötC cells using saporin-Substance P progressively disrupts breathing and eventually stops it (Gray *et al*., 2001), with this effect more pronounced during sleep (McKay & Feldman, 2008). Substance P expression is also consistent with the immunohistochemical distribution of *Dbx1* neurons in the preBötC (Sun *et al*., 2019). Thus, substance P and its cognate NK-1Rs are likely excitatory and critical for the generation of breathing. In the present study, activation of *Tac1*-expressing preBötC cells promotes inspiration to similar levels observed when glutamatergic preBötC cells were stimulated. According to *in situ* hybridization, *Tac1*-expressing cells constitute less than 24% of glutamatergic preBötC cells. Consistent with our results, stimulation of 4 to 9 glutamatergic neurons in brainstem slice initiated an inspiratory burst with levels comparable to endogenous burst (Kam *et al*., 2013). Overall, *Tac1* cells constitute a population of preBötC cells sufficient to generate inspiratory activity *in vivo*.

### Phase-dependent role of *Tac1*-expressing cells

Photostimulation of *Tac1*-expressing preBötC cells produced inspiratory activity only during discrete periods of the respiratory cycle. *Tac1* photostimulation had no effect during the first 20% of the respiratory cycle but triggered inspiratory activity when applied during post-inspiration or expiration. These results are consistent with the lack of effects of *Vglut2* or *Dbx1* stimulation during the inspiratory phase in anesthetized mice (Oliveira et al., 2021). Consistent with our results *Vglut2*, but not *Dbx1*, stimulation produced inspiration when photostimulation was applied during expiration (Oliveira *et al*., 2021). On the other hand, stimulation of *Dbx1* neurons in brainstem slides *in vitro* produced inspiration two seconds after endogenous inspiratory bursts (Kottick and Del Negro, 2015). A smaller and more specific population of excitatory SST-expressing cells (87% glutamatergic) also showed excitatory effects, where it prolonged cycle duration in anesthetized mice but shortened cycle duration when stimulation was performed during mid-expiration (de Sousa Abreu *et al*., 2022). In this study, the resistance of preBötC cells to initiate another inspiratory burst during the early phase of respiratory cycle (0-20%) suggests that *Tac1* cells, like other rhythmogenic glutamatergic cells, undergo a refractory period. This prolonged period of reduced network excitability after an inspiratory burst (Baertsch *et al*., 2018) may be due to prolonged afterhyperpolarization following inspiration or depletion of presynaptic vesicles (Ramirez and Baertsch, 2018).

### *Tac1* preBötC cells and motor behaviors

The preBötC is the site of respiratory rhythmogenesis and is critical to generate inspiration in mammals. The preBötC is not limited to its main role in the generation of breathing. A subset of preBötC cells expressing *Dbx1* and cadherin are also involved in arousal, and their ablation increased slow-wake cortical activity (Yackle *et al*., 2017), suggesting that the preBötC may play roles beyond rhythmic breathing. Outside of this region, activation of *Tac1* neurons in the preoptic area of the hypothalamus (POA) promoted arousal and enhanced locomotor activity (Reitz et al., 2021). Substance P is also a key-neuropeptide involved in nociception (Mantyh, 2002) and locomotion (Farrell *et al*., 2021). Descending *Tac1* brainstem circuits mediate behavioral responses, such as the fight-or-flight response, associated with brisk locomotor activity (Barik *et al*., 2018; Kuwaki, 2021). To anticipate the body’s metabolic demand in the event of locomotor nocifensive response, nociceptive stimuli elicit cardio-respiratory responses, such as augmented breathing (Jafari *et al*., 2017) and cardiovascular response in conscious rodents (Unger et al., 1988). In our study, stimulation of *Tac1* cells promoted locomotion or movement. It is plausible that the locomotor response to photostimulation of *Tac1* preBötC cells may be due to the photostimulation of adjacent motor nuclei. Most of ChETA expression (shown by *Eyfp* mRNA) was restricted to the preBötC, the caudal ventrolateral reticular nucleus (CVL), and the Ventral SpinoCerebellar tract (VSC), with moderate expression in the Lateral ParaGigantocellular nucleus (LPGi) (**Suppl. Figure 3**). The CVL is involved in nociceptive-cardiovascular integration (Lima *et al*., 2002) suggesting that this nucleus may not contribute directly to increased locomotor activity observed with *Tac1* photostimulation. Although the VSC drives maintenance and control of locomotion in rodents (Chalif *et al*., 2022) and the LPGi controls locomotion (Capelli *et al*., 2017), their locations medial to the optical fiber suggest that they were unlikely photostimulated by laser light. In addition, the LPGi did not present strong expression of *Tac1* mRNAs. Although it cannot be excluded that some motor nuclei may be partially stimulated by light, it is unlikely that activation of a few motor cells mediate the strong locomotor response observed with *Tac1* photostimulation. Here, we conclude that *Tac1*-expressing preBötC neurons constitute a population of neurons linking breathing and locomotion.

### Reversal of respiratory depression by opioid drugs

Opioid drugs present unwanted side effects such as respiratory depression. The respiratory properties of opioid drugs are due to their action on µ-opioid receptors (Heinricher *et al*., 2009; Montandon, 2022). The preBötC expresses MORs and mediates a large component of respiratory rate depression by opioid drugs (Montandon *et al*., 2011; Bachmutsky *et al*., 2020; Varga *et al*., 2020). Interestingly, preBötC neurons expressing NK-1R, the cognate receptors of substance P, are preferentially inhibited by opioid drugs (Montandon *et al*., 2011). In our study, we found co-expression of *Oprm1* (the gene coding for MORs) and *Tac1* in preBötC neurons, and stimulation of *Tac1* preBötC cells entirely reversed respiratory depression by fentanyl. Consistent with these results, opioid drugs directly inhibit substance P-expressing cells and substance P production in the spinal cord (Fukazawa *et al*., 2007). Moreover, an excitatory *Tac1* spinal-brainstem circuit mediates noxious responses in rodents (Barik *et al*., 2018; Gutierrez *et al*., 2019) and deletion of the *Tac1* gene in mice increases the inhibitory effects of opioid drugs, suggesting that substance P could reverse respiratory depression by opioid drugs (Takita *et al*., 2000; Berner *et al*., 2012). The tachykinin system (substance P release and activation of NK-1R) may therefore constitute a tonic excitatory circuit that could antagonize the suppressive effects of opioids. Consistent with this hypothesis, we showed that stimulation of *Tac1* preBötC cells alleviated respiratory depression by fentanyl in intact animals. Interestingly, stimulation of breathing by NK-1R activation in the preBötC is regulated by G-protein-gated inwardly rectifying potassium channels (GIRK) (Montandon *et al*., 2016a), which also regulates respiratory rate depression by opioid drugs (Montandon *et al*., 2016b). Such converging cellular mechanisms further support the idea of a link between *Tac1* neurons and preBötC inhibition by drugs acting on MORs.

The brainstem circuits generating and regulating rhythmic breathing are complex. At the core of the respiratory circuits is the preBötC, a collection of neurons with diverse neurochemical identities. Here, we identified a sub-population of glutamatergic neurons, expressing the precursor *Tac1* of the neuropeptide substance P, that can trigger inspiration and entrain breathing in freely-behaving mice. The importance of *Tac1* preBötC neurons is highlighted by the fact that it can produce inspiration when breathing is inhibited by opioid drugs. Importantly, activation of these neurons also triggers locomotion therefore suggesting that *Tac1* preBötC neurons may play roles beyond the generation of rhythmic breathing in freely-moving mice.

---

**Supplementary Figure 1.**
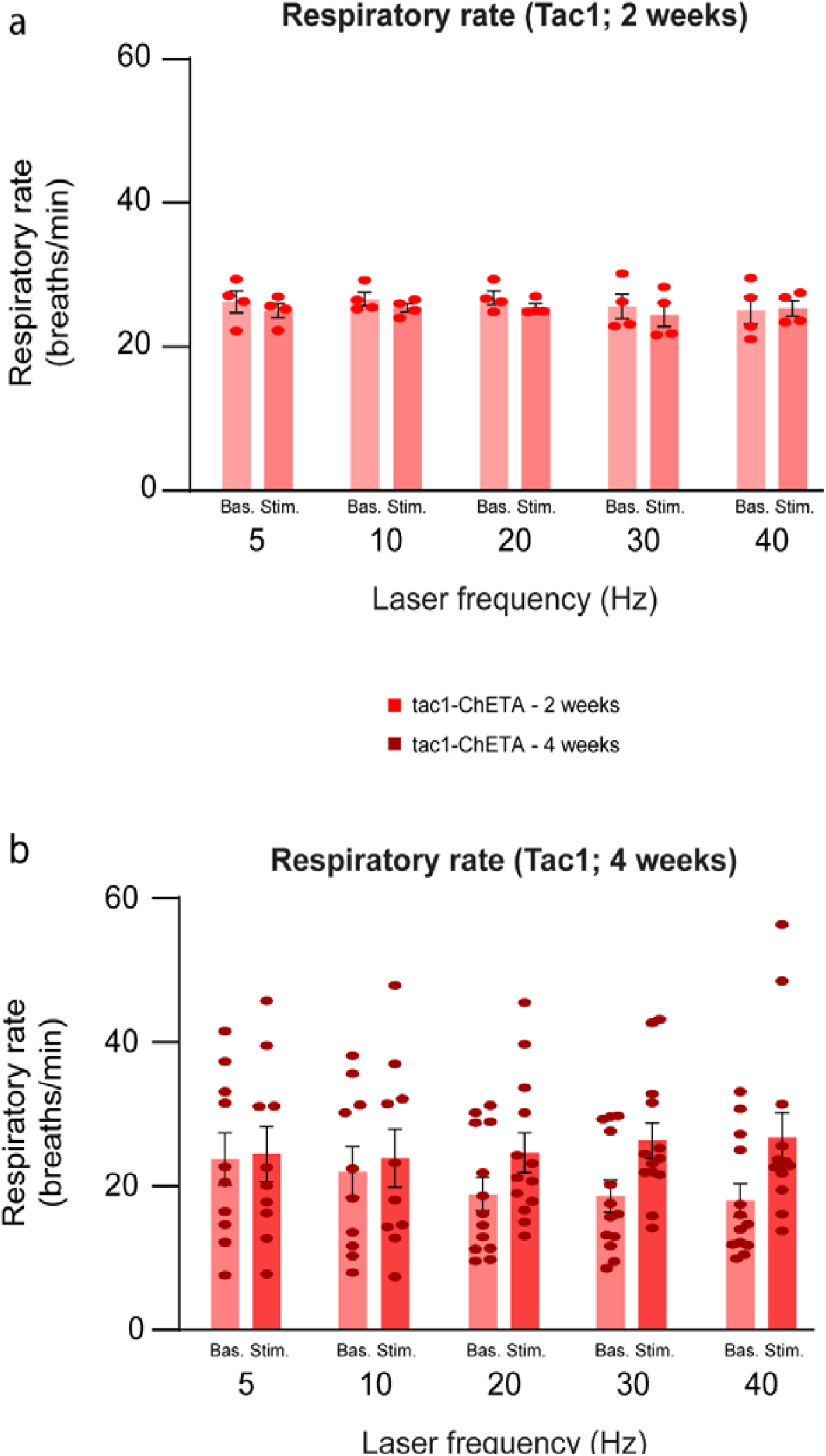
Optogenetic stimulation of *Tac1* preBötC cells in anesthetized mice. **(a)** Mean diaphragm respiratory rate (breaths/min) before (Baseline) and during (Stimulation) laser stimulation in Tac1-ChETA mice 2 weeks post virus injection (laser stimulation effect: *P=0.4403*, n=8). Laser stimulations were performed at 5, 10, 20, 30 and 40 Hz. **(b)** Respiratory rate were also measured in *Tac1*-ChETA mice 4 weeks post virus injection (laser stimulation effect: *P=0.1497*, n=30). Laser stimulations were performed at 5, 10, 20, 30 and 40 Hz. Data are presented as means ± SEM, with individual data points.

**Supplementary Figure 2.**
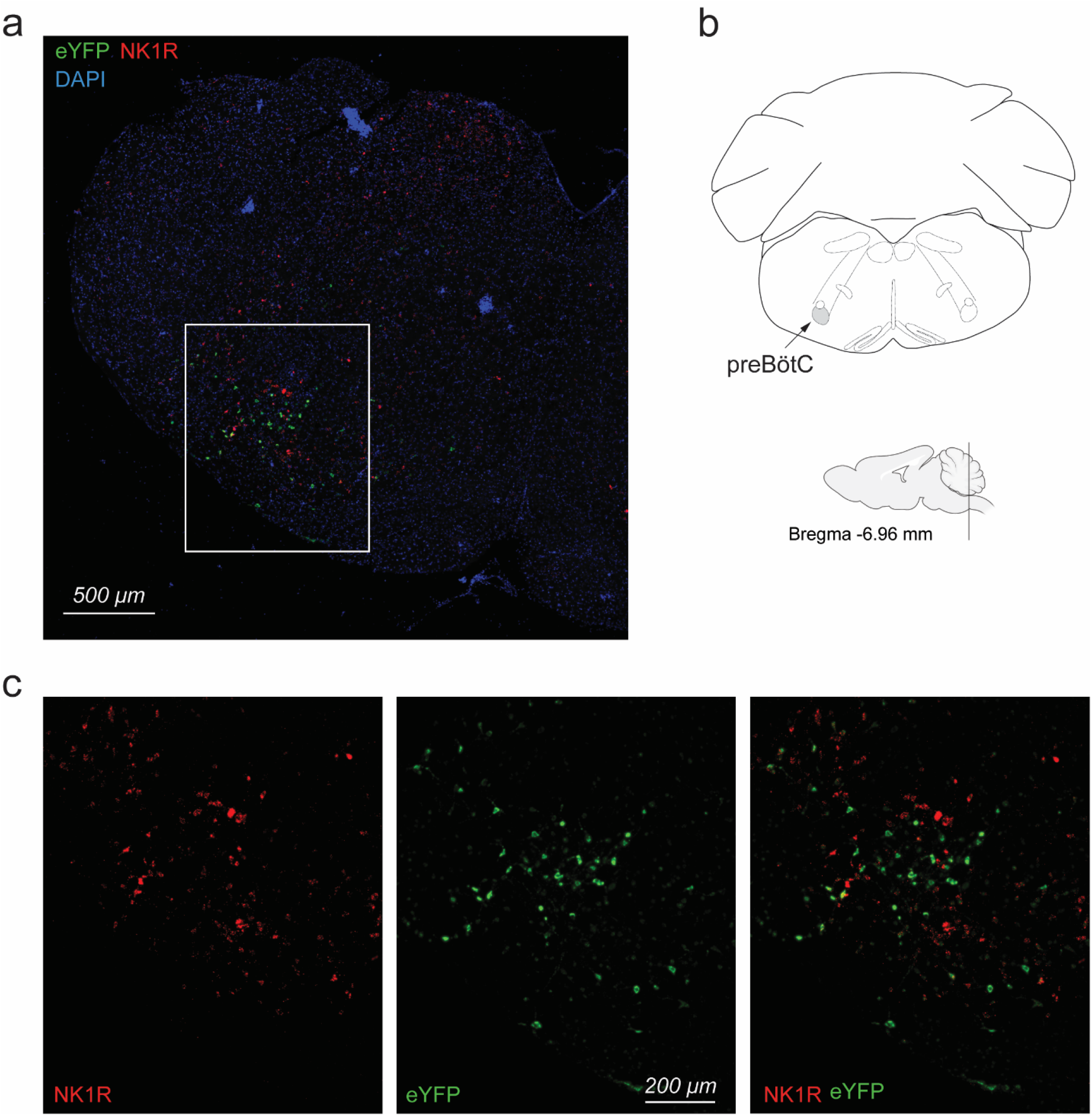
*In-situ* hybridization showing mRNA expression for eYFP (ChETA) in the preBötC region shown by neurokinin-1 receptors (*Tacr1* or NK-1R) in *Tac1-*cre mice. **(a)** Injection of the adenoassociated virus carrying ChETA in the preBötC was confirmed using *in-situ* hybridization. eYFP (green) mRNA expression is located in the same region than expression of NK-1R (red, encoded by the gene *Tacr1*). **(b)** The preBötC region is well demarcated by NK-1R expression. (**c)** Magnified image of the preBötC region showing mRNA expression of eYFP (for ChETA, green) and *Tacr1* (for NK1R, red). DAPI is shown in blue. Note that although ChETA (green) is co-expressed with *Tac1*, it is not necessarily c-expressed with neurokinin-1 receptor cells (or *Tacr1*, red), but located in the same region.

**Supplementary Figure 3.**
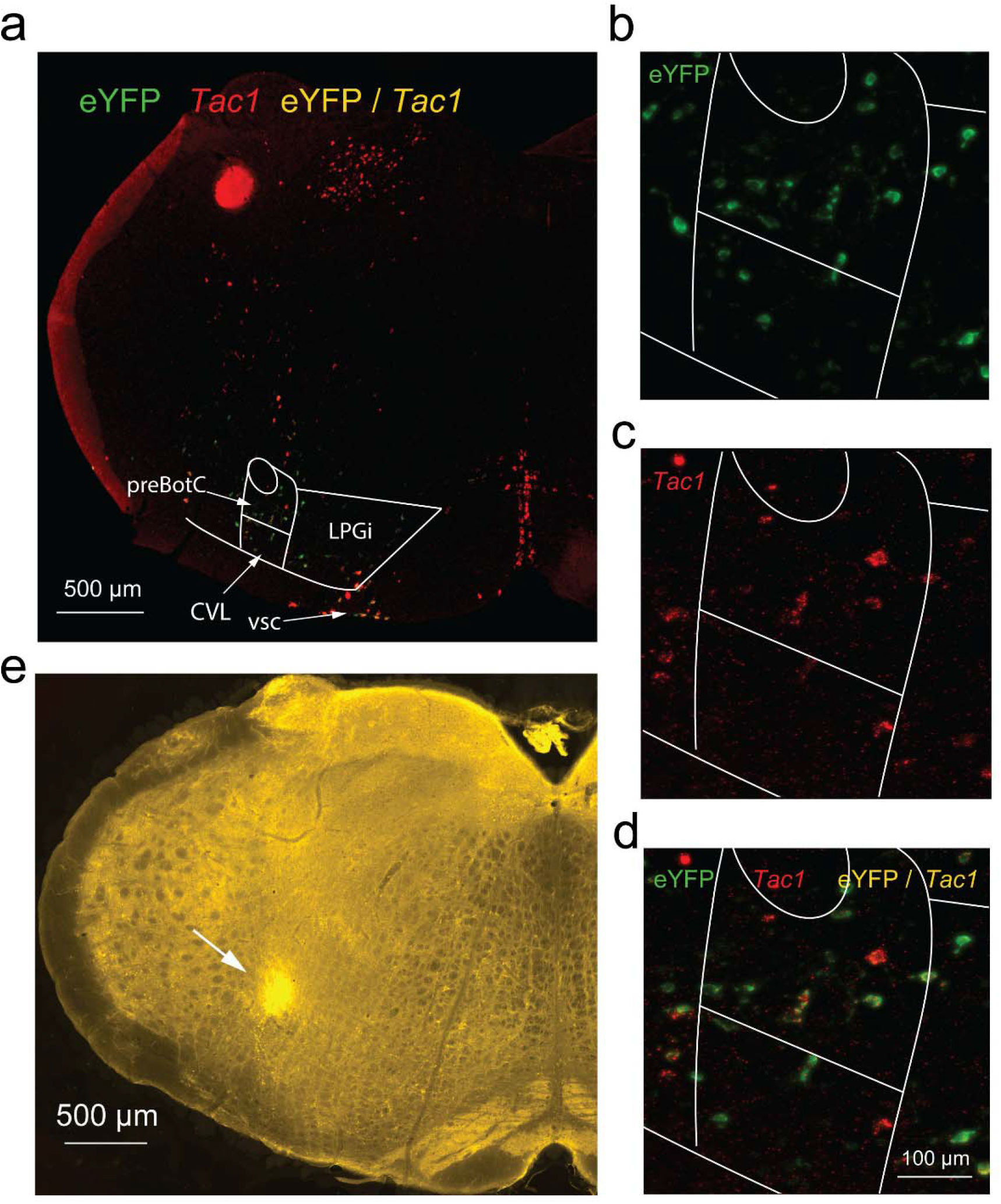
mRNA expressions of *eYFP* (marking ChETA) and *Tac1* (substance P) in the preBötzinger Complex region and adjacent motor nuclei. **(a)** Using the Paxinos’s mouse atlas, we delineated the motor nuclei surrounding the preBötC. Ventral to the preBötC is located the Caudal VentroLateral reticular nucleus (CVL). Medial to the preBötC is located the Lateral ParaGigantocellular nucleus (LPGi). Ventral to the the LPGi is located the Ventral SpinoCerebellar tract (VSC). **(b)** In this experiment, eYFP (ChETA) was found in the preBötC, the CVL, the LPGi and the VSC. **(c)** Tac1 expression was observed in the preBötC and VSC, but only moderately in the CVL and LPGi. **(d)** In fact most of co-expression of Tac1 and eYFP was observed in the preBötC and the VSC. **(e)** The optical fiber was positioned dorsal to the preBötC and lateral to the LPGi and the VSC.

**Supplementary Figure 4.**
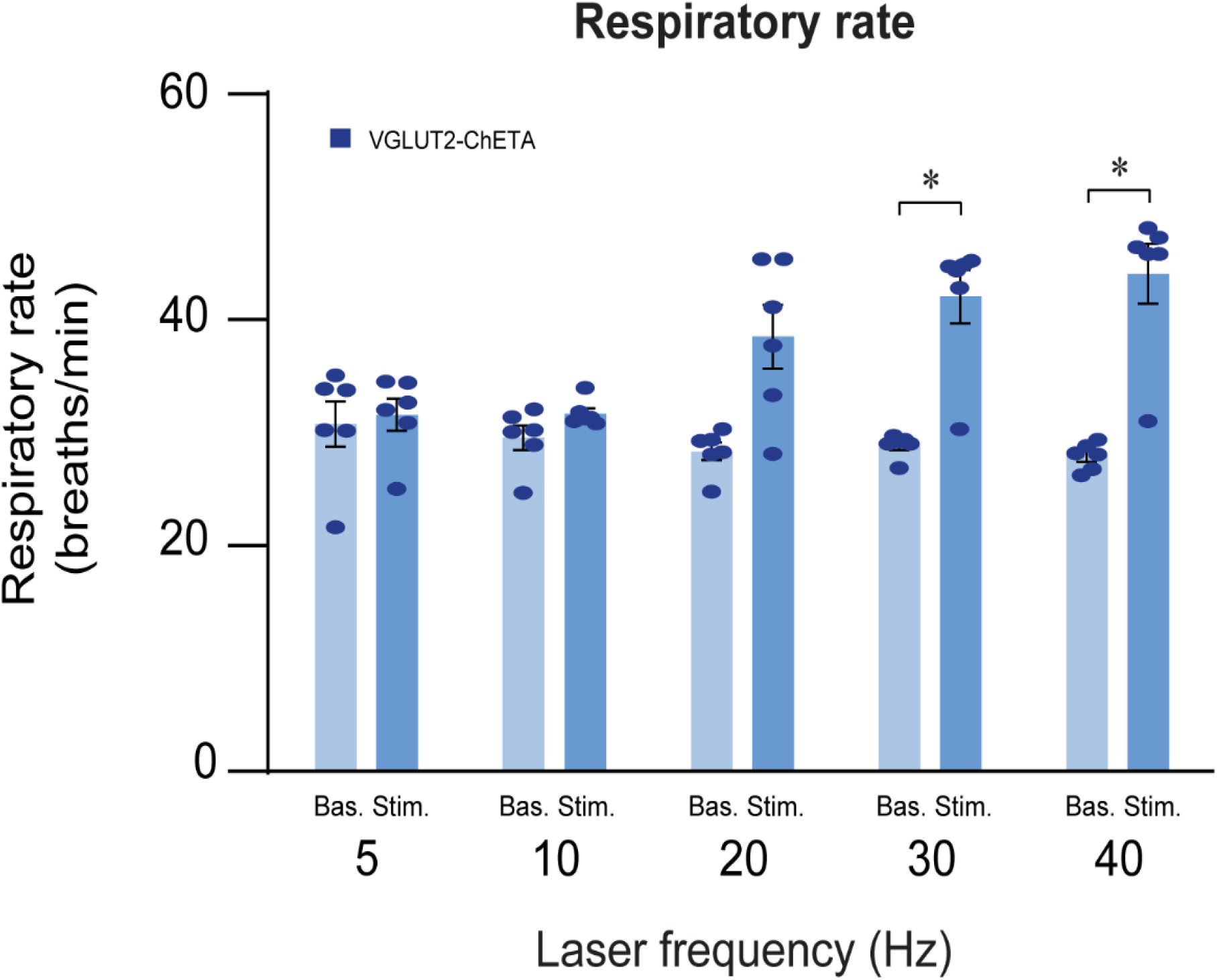
Optogenetic stimulation of *Vglut2* preBötC cells in anesthetized animals. Mean diaphragm respiratory rate (breaths/min) before (Baseline) and during (Stimulation) laser stimulation in Vglut2-ChETA mice 2 weeks post virus injection (laser stimulation effect: *P=0.0007*, n=12). Laser stimulations were performed at 5, 10, 20, 30 and 40 Hz. Data are presented as means ± SEM, with individual data points. * indicate means significantly different from corresponding baseline value with P<0.05.

**Supplementary Figure 5.**
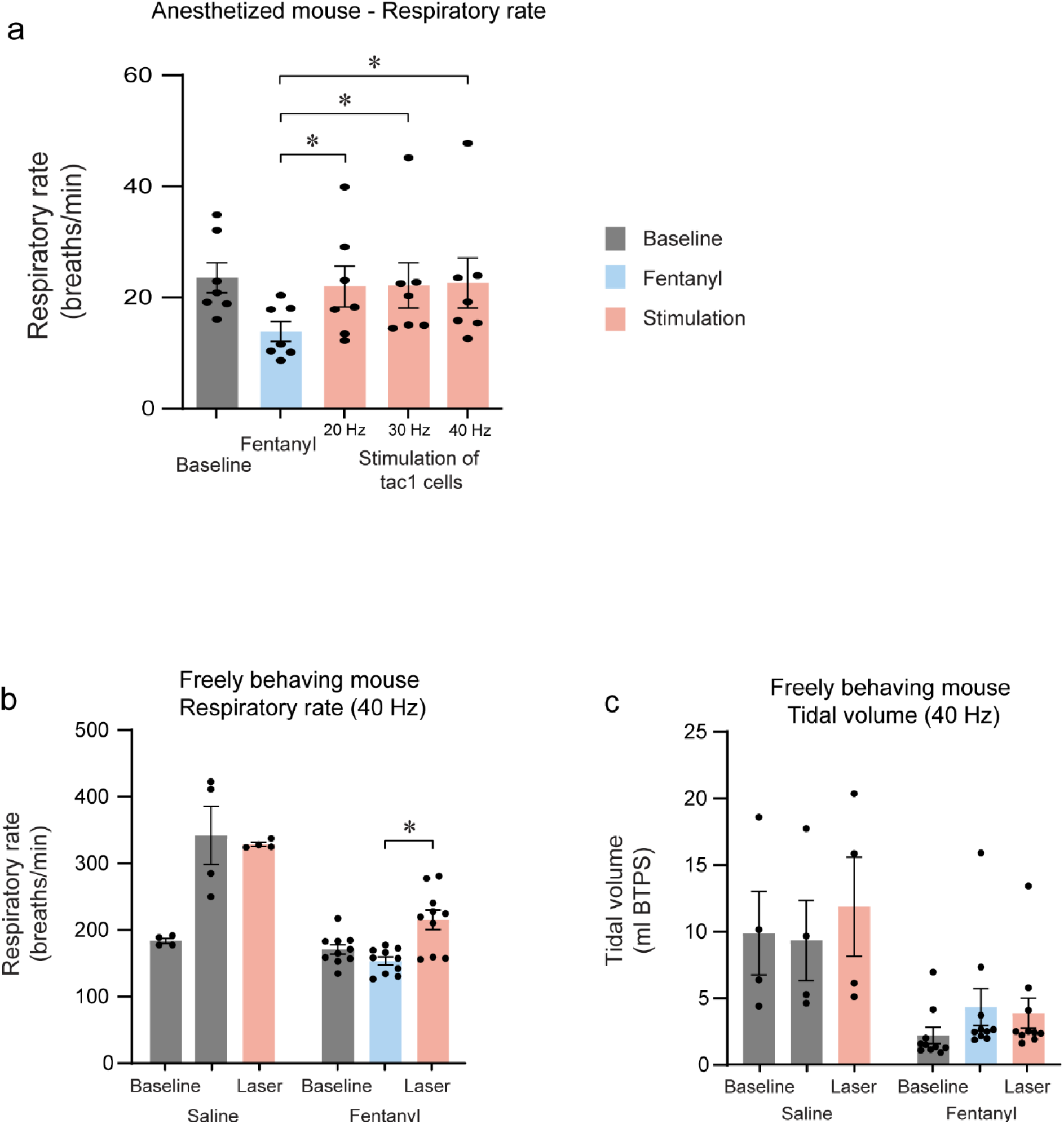
Fentanyl depression and optogenetic stimulation of *Tac1* preBötC cells. **(a)** Mean diaphragm respiratory rate (breaths/min) following intramuscular injection of fentanyl (5µg/kg) and different laser stimulations (20, 30, 40 Hz) in anesthetized Tac1-ChETA mice 4 weeks post virus injection. (laser stimulation effect: *P=0.0008*, n=7). **(b)** Mean respiratory rate (breaths/min) (laser stimulation effect: *P<0.0001*, n=14) and **(c)** mean tidal volume (laser stimulation effect: *P=0.0174*, n=14) assessed using whole-body plethysmography, following intraperitoneal injection of fentanyl (0.3 mg/Kg) or saline and laser stimulation (40 Hz) in freely-moving, non-anesthetized, *Tac1-*ChETA mice 4 weeks post virus injection. Data are presented as means ± SEM, with individual data points. * indicate means significantly different from corresponding fentanyl value with P<0.05.

